# *Enterococcus* peptidoglycan remodeling promotes immune checkpoint inhibitor therapy

**DOI:** 10.1101/2020.08.20.256263

**Authors:** Matthew E. Griffin, Juliel Espinosa, Jessica L. Becker, Jyoti K. Jha, Gary R. Fanger, Howard C. Hang

## Abstract

The antitumor efficacy of cancer immunotherapy has been correlated with specific species within the gut microbiota. However, molecular mechanisms by which these microbes affect host response to immunotherapy remain elusive. Here we show that specific members of the bacterial genus *Enterococcus* can promote anti-PD-L1 immunotherapy in mouse tumor models. The active enterococci express and secrete orthologs of the NlpC/p60 peptidoglycan hydrolase SagA that generate immune-active muropeptides. Expression of SagA in non-protective *E. faecalis* was sufficient to promote antitumor activity of clinically approved checkpoint targets, and its activity required the peptidoglycan sensor Nod2. Notably, SagA-engineered probiotics or synthetic muropeptides also promoted checkpoint inhibitor antitumor activity. Our data suggest that microbiota species with unique peptidoglycan remodeling activity may enhance immunotherapy and could be leveraged for next-generation adjuvants.

**One Sentence Summary:** A conserved family of secreted NlpC/p60 peptidoglycan hydrolases from *Enterococcus* promote antitumor activity of immune checkpoint inhibitors.

## Main Text

Cancer immunotherapy harnesses host immune mechanisms to impede tumor growth and has demonstrated clinical success across a range of solid and liquid tumors (*1–3*). In particular, antibodies that target immune inhibitory pathways such as CTLA-4 and the PD-1/PD-L1 signaling axis have been approved to treat a wide array of human cancers (*3*). However, patient response to immune checkpoint inhibitors is variable (*3*). The success of checkpoint blockade relies on numerous factors including mutational burden of the malignancy (*4*), successful tumor antigen presentation (*5*), recruitment and infiltration of lymphocytes (*6*), and signaling cues within the tumor microenvironment (*7*). Recently, the gut microbiota has emerged as a potent new factor associated with the efficacy of anti–CTLA4, anti–PD-1, and anti–PD-L1 treatment (*8–13*). In animal and human cohorts, the presence of specific microbial species was correlated with responsiveness to checkpoint blockade agents. The antitumor activity of these microbes was recapitulated in preclinical mouse models upon co-housing, fecal transplant, or direct inoculation, suggesting that the correlated microorganisms are direct causative agents of improved therapeutic response. Nevertheless, little is known about the exact, molecular factors by which these immune modulatory microbes act.

Multiple analyses of the commensal microbiome from human cohorts treated with immunotherapies targeting PD-1 have revealed that the bacterial genus *Enterococcus* is enriched in responding patients (*12, 13*). Although antibiotic-resistant strains of *E. faecium* and *E. faecalis* can be pathogenic (*14*), commensal strains of these bacteria have been used employed as probiotics in animals and humans (*15*). Recent studies also suggested that *Enterococcus* species can trigger immune signaling pathways and modulate infection (*16–18*), autoimmunity (*19*), and graft-versus-host-disease (*20*). These observations prompted our inquiry into whether specific *Enterococcus* species and strains are sufficient to improve checkpoint blockade.

To evaluate specific enterococci and their mechanism of action, we utilized mouse tumor models and orally administered bacteria. Here, specific pathogen-free (SPF) C57BL/6 mice from The Jackson Laboratory were pretreated for two weeks with a broad-spectrum antibiotic cocktail to clear resident microbial species and subsequently provided water supplemented with bacteria. Supplemented animals were then subcutaneously implanted with B16-F10 melanoma cells and treated with anti–PD-L1 (**Fig. 1A**). We first focused on the two most common *Enterococcus* species in the human gut microbiota: *E. faecium* (Efm) and *E. faecalis* (Efs). We found that supplementation with the human commensal *E. faecium* strain Com15 without therapeutic intervention did not alter tumor growth (**Fig. 1B**). However, treatment of supplemented animals with anti–PD-L1 at both high and low dosages showed a significant decrease in tumor size compared to mice treated with anti–PD-L1 alone. Using the low anti–PD-L1 dose protocol going forward, we then compared the activity of *E. faecium* with *E. faecalis* across multiple strains of each species—human-isolated, type, and multi-drug resistant—to ascertain if the observed synergistic activity was unique to *E. faecium* Com15. Inhibition of tumor growth was observed across all three isolates of *E. faecium* and not for any strain of *E. faecalis* (**Fig. 1C**). To ensure that these effects were not due to differences in bacterial load, fecal samples were plated onto *Enterococcus*-selective medium and enumerated. All bacterial strains yielded similar amounts of colony forming units (CFU, **Fig. 1D**), indicating that the observed antitumor activity of *E. faecium* strains was not due to differences in bacterial amount in the gut. In addition to *E. faecium*, 16S rRNA analysis of gut microbiota from responsive human patients also uncovered the enrichment of other *Enterococcus* species (*12*). We found that antitumor activity was conserved across multiple species including *E. durans* (Eds), *E. hirae* (Ehe), and *E. mundtii* (Emi) (**Fig. 1E and S1A**). CFU analysis of the colonized animals again revealed that activity did not correlate with bacterial load (**Fig. 1F and S1B**).

**Fig. 1.**
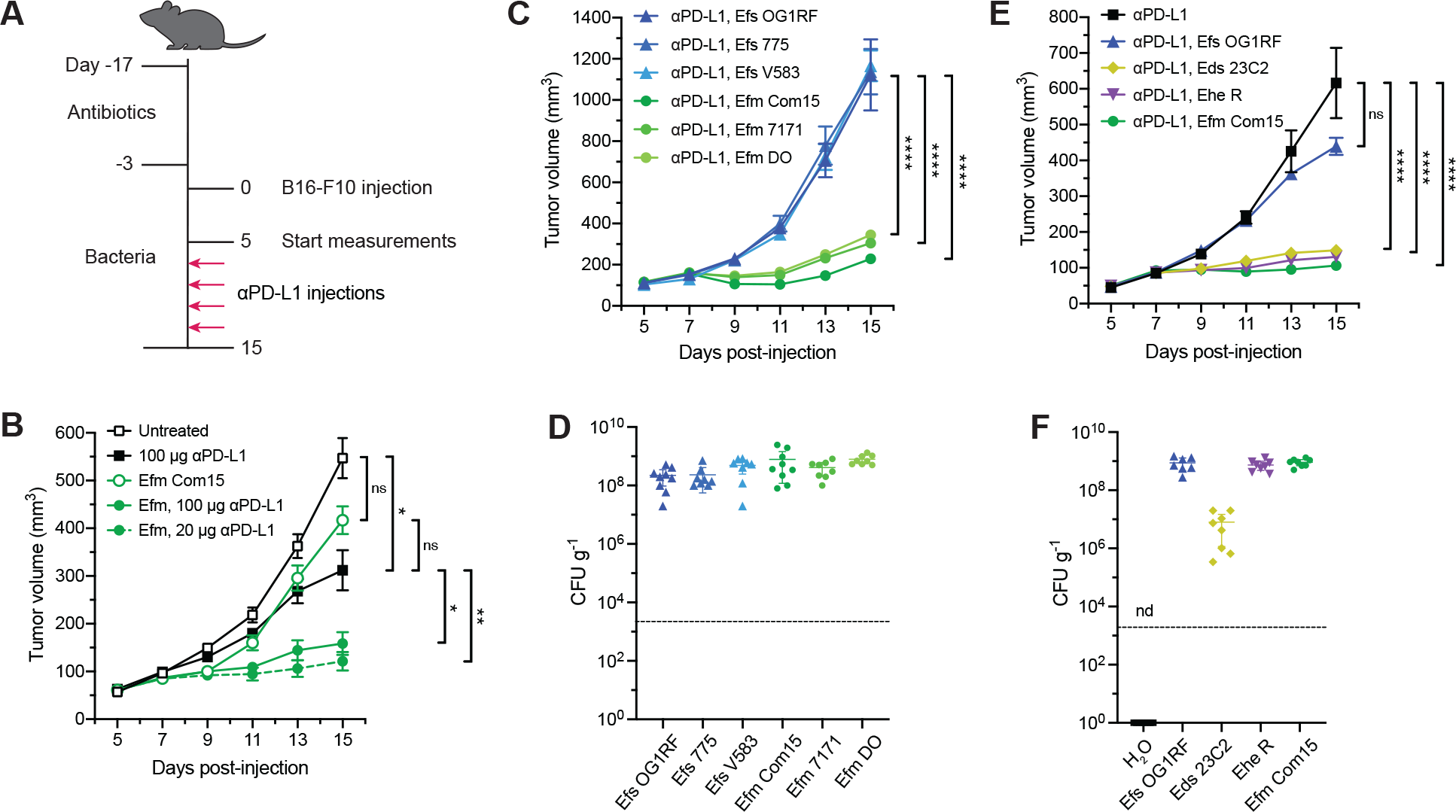
Specific enterococci improve anti–PD-L1 efficacy in B16-F10 melanoma model. (**A**) Schematic of tumor growth model in SPF mice with antibiotic treatment and oral enterococci supplementation. Days are indexed based on the day of tumor injection. Mice were provided antibiotic-containing water *ab libitum* for two weeks followed by water supplemented with the indicated enterococci for the remainder of the experiment. Animals were then subcutaneously implanted with B16-F10 melanoma cells, and tumor volume measurements started when tumors reached ~50-100 cm^3^ (day 5). Mice were treated with anti–PD-L1 by intraperitoneal injection every other day starting two days after the start of measurement. For all data except for (B), 20 μg anti–PD-L1 was used for each injection. (**B**) B16-F10 tumor growth in antibiotic-treated animals that were supplemented with or without *E. faecium* (Efm) Com15 and treated with or without anti–PD-L1 starting on day 7 at doses indicated in the legend. *n* = 7-8 mice per group. (**C**) B16-F10 tumor growth in antibiotic-treated mice that were supplemented with the indicated *E. faecalis* (Efs) and Efm strains and treated with anti–-PD-L1 starting on day 7. *n* = 7-8 mice per group. (**D**) Colony forming unit (CFU) analysis of Efs and Efm strains in fecal samples harvested from animals as treated in (C). (**E**) B16-F10 tumor growth in antibiotic-treated mice that were supplemented with the indicated enterococci and treated with anti–PD-L1 starting in day 7. *n* = 8-9 mice per group. (**F**) CFU analysis of enterococci in fecal samples harvested from animals as treated in (E). nd = not detected. For (B), (C), and (E), data represent as mean ± s.e.m. and were analyzed by mixed-effects model with Tukey’s multiple comparisons post-hoc test. \**P* < 0.05, \*\**P* < 0.01, \*\*\*\**P* < 0.0001, ns = not significant. For (D) and (F), each symbol represents one mouse. Dotted lines indicate the limit of detection (2,000 CFU g^-1^). Data represent means ± 95% confidence interval.

To explain the species-specific differences we observed across the *Enterococcus* genus, we examined possible sources of their immunomodulatory activity. Notably, our previous work indicated that *E. faecium* has unique peptidoglycan composition and remodeling capabilities to enhance host tolerance to enteric pathogens (*16–18*). To compare peptidoglycan composition across the enterococci, we isolated sacculus of each species and analyzed the digested peptidoglycan fragments by high performance liquid chromatography–mass spectrometry (HPLC-MS). All four of the immunotherapy-active enterococci (*E. faecium, E. durans, E. hirae,* and *E. mundtii*) showed similar peptidoglycan fragment patterns compared to the nonactive species *E. faecalis* and *E. gallinarum* (Egm) (**Fig. 2A and S2A**). Additionally, all four immunotherapy-active species showed an abundance of smaller, non-crosslinked peptidoglycan fragments, suggesting higher levels of peptidoglycan remodeling and turnover. The modification of peptidoglycan stem peptides is catalyzed by conserved families of amidases and peptidases (*21*), so we examined *Enterococcus* genomic assemblies for unique expression of peptidoglycan remodeling enzymes (**Fig. S3**). Based on this analysis, we identified a cluster of NlpC/p60 hydrolases that were highly conserved through all active enterococci (**Fig. 2B**). This group of related enzymes contained the peptidoglycan hydrolase secreted antigen A (SagA) from *E. faecium,* which we previously demonstrated improves host immunity against enteric infections (*16–18*). Using primary sequence alignment, we found that the putative SagA orthologs from other active enterococci were highly homologous, with greater than 90% sequence identity within the C-terminal NlpC/p60 hydrolase domain (**Fig. 2C**). Conversely, the closest related protein in *E. gallinarum* showed only 67% sequence identity in the predicted catalytic domain (**Fig. S2b**). No direct orthologs were found in *E. faecalis,* and the two most similar conserved enzymes, *sagA-like* proteins A and B (SalA and SalB) (*22*), possessed distinct C-terminal domains (**Fig. 2C**). Structural modeling revealed that the putative SagA orthologs also shared similar secondary structural organization compared to a crystal structure of the NlpC/p60 hydrolase domain from *E. faecium* SagA (**Fig. S4**).

**Fig. 2.**
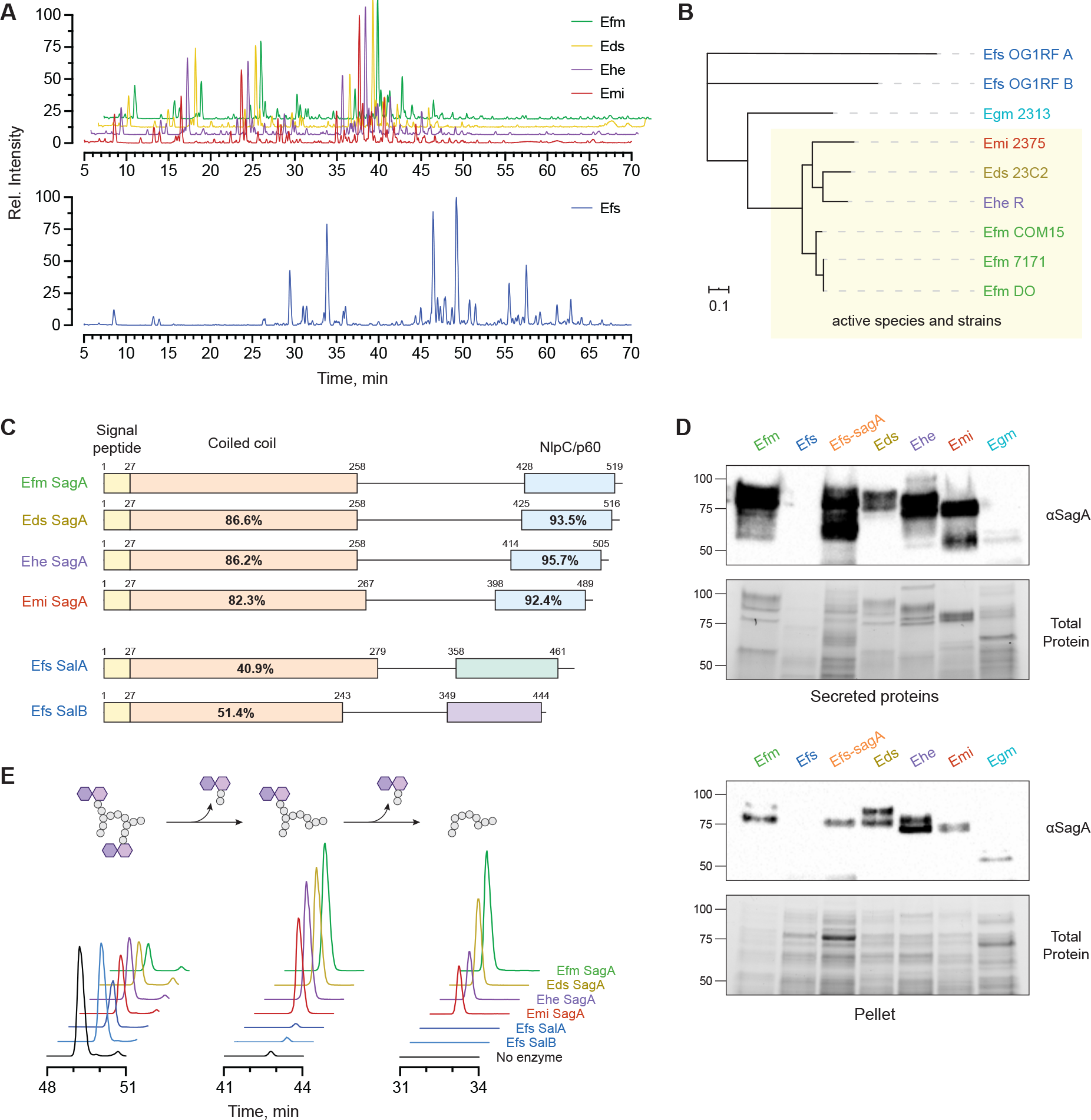
Protective enterococci express and secrete active orthologs of the peptidoglycan NlpC/p60 hydrolase SagA. (**A**) LC-MS ion chromatograms of mutanolysin-treated peptidoglycan harvested from *Enterococcus* species. Data are shown as relative intensity of base peak ion abundance. (**B**) Unrooted phylogenetic clustering of putative SagA ortholog protein sequences identified by global peptidoglycan peptidase analysis of enterococci species and strains along with the closest entries from Egm and Efs based on IQ-Tree analysis. Active strains are indicated by the yellow box. Scale bar represents sequence distance. (**C**) Comparison of primary sequence homology and domain architecture of SagA orthologs and two conserved *sagA-like* proteins from Efs. Numbers above each bar denote amino acid residue boundaries of the indicated domains, and percentages represent sequence identity compared to Efm SagA. (**D**) Western blot detection of SagA orthologs in secreted protein and cell pellet fractions harvested from overnight cultures of the indicated enterococci using antiserum raised against Efm Com15 SagA. Bottom panels show total protein loading. Numbers indicate estimated molecular weight (kDa). (**E**) *In vitro* activity of purified SagA orthologs. Data are shown as extracted LC-MS ion chromatograms of a crosslinked peptidoglycan substrate and two iterative hydrolysis products after incubating a mixture of peptidoglycan fragments with purified SagA orthologs from the indicated species for 16 hours at 37 °C. Peak heights are shown as relative intensity of ion abundance, and all chromatograms are shown on the same scale.

To directly detect the expression of SagA orthologs in enterococci, we performed Western blotting on both the secreted and cell-associated protein fractions from each species. Using antiserum raised against *E. faecium* SagA (*18*), strong signals were observed in the supernatant fractions of *E. faecium, E. durans, E. hirae,* and *E. mundtii* but not *E. faecalis* or *E. gallinarum* (**Fig. 2D**), suggesting that the similar proteins are both highly expressed and secreted by these species. The signal was confirmed to be SagA-dependent using a strain of *E. faecalis* OG1RF engineered with a chromosomal *sagA* insertion (*Efs-sagA*). As expected, all tested strains of *E. faecium* also showed similar expression and secretion patterns for SagA (**Fig. S5**). To determine if these SagA orthologs were functional, we analyzed the hydrolytic activity of the purified recombinant proteins on peptidoglycan *in vitro* (**Fig. 2E and S6**). Proteins from *E. durans, E. hirae,* and *E. mundtii* showed D,L-endopeptidase activity against a model crosslinked peptidoglycan fragment similar to *E. faecium* SagA (**Fig. S7**). The hydrolytic activity of SagA was confirmed using a mutant construct lacking the cysteine active site residue, which was conserved in all orthologs (**Fig. S8**). *E. faecalis* SalB did not hydrolyze peptidoglycan at detectable levels in our assay, whereas SalA cleaved the crosslinked fragment in the cross-bridge region rather than the peptide stem (**Fig. 2E and S9**). These results show the immunotherapy-active enterococci possess similar peptidoglycan composition and remodeling activity.

We then investigated whether SagA was sufficient to enhance the efficacy of checkpoint inhibitor therapy. Since *sagA* is an essential gene in *E. faecium* (*23*), we compared the inactive, parental *E. faecalis* OG1RF strain with the engineered, SagA-expressing strain *E. faecalis–sagA* (**Fig. 2D**). The peptidoglycan profile of *E. faecalis–sagA* showed changes consistent with increased D,L-endopeptidase activity (**Fig. S10A**), indicative of active SagA expression (**Fig. 2D**). Antibiotic-treated animals that were orally supplemented with *E. faecalis–sagA* showed a significant decrease in B16-F10 tumor growth upon anti–PD-L1 compared to the parental *E. faecalis* strain (**Fig. 3A**). This antitumor activity was similar to the phenotype observed in *E. faecium*-supplemented animals. Furthermore, fecal CFU analysis showed that *E. faecalis–sagA* exhibited similar bacterial load as the parental *E. faecalis* strain and *E. faecium* (**Fig. S10B**). We then tested the outcome of SagA expression on checkpoint inhibitor therapy in an animal model with an intact, complex microbiota. SPF C57BL/6 mice from Taconic Biosciences were chosen as previous reports have found that these mice do not respond strongly to anti–PD-L1 therapy similar to germ-free animals (*9*, *13*). Here, these mice were directly supplemented with enterococci-containing water without antibiotic pre-treatment prior to B16-F10 implantation (**Fig. S11A**). As seen in the antibiotic-pretreated model, animals supplemented with *E. faecium* or *E. faecalis–sagA* showed a significant decrease in tumor growth compared to treatment with antibody alone or antibody and parental *E. faecalis* (**Fig. S11B**). We found much lower overall *Enterococcus* CFU counts that did not significantly differ between the supplemented mice, suggesting that this effect did not require enteric domination by the newly administered, active enterococci (**Fig. S11C**). We then asked whether the antitumor phenotype occurred through changes in endogenous microbiota populations caused by SagA-expressing enterococci. Fecal samples were collected prior to and during bacterial administration and compared by 16S rRNA sequencing. No significant differences in microbiota composition were observed between samples regardless of species administered or duration (**Fig. S11, D-G**).

**Fig. 3.**
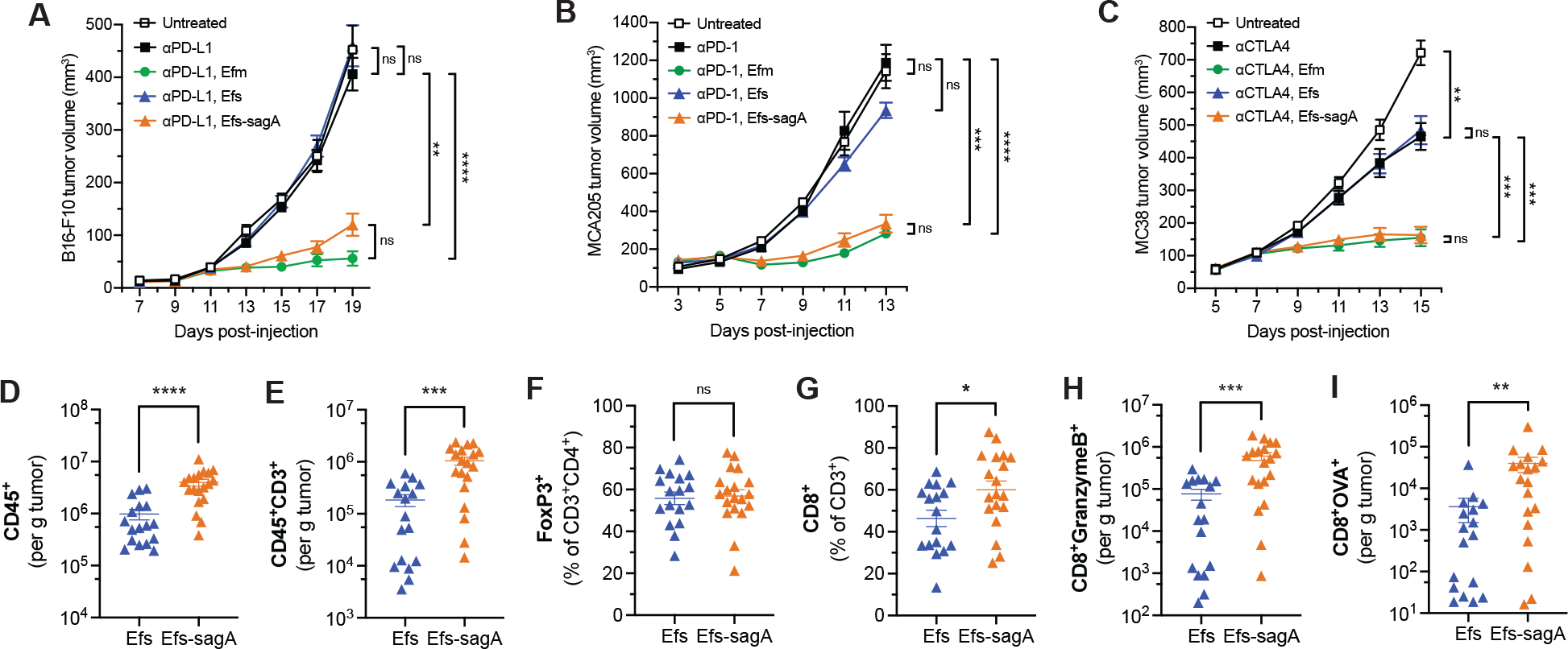
SagA is sufficient to broadly improve immune checkpoint inhibitors and elicit an adaptive immune response. (**A**) B16-F10 tumor growth in antibiotic-treated mice that were supplemented with the indicated enterococci and treated with anti–PD-L1 starting on day 9. *n* = 8 mice per group. (**B**) MCA205 tumor growth in antibiotic-treated mice that were supplemented with the indicated enterococci and treated with anti–PD-1 starting on day 5. *n* = 8 mice per group. (**C**) MC38 tumor growth in antibiotic-treated mice that were supplemented with the indicated enterococci and treated with anti–CTLA-4 starting on day 7. *n* = 8 mice per group. Data for (A)-(C) represent means ± s.e.m. and analyzed by mixed-effects model with Tukey’s multiple comparisons post-hoc test. (**D**) to (**H**) Quantification of tumor infiltrating CD45^+^ cells (D), total CD3^+^ T cells (E), FoxP3^+^ regulatory T cells (F), CD8^+^ T cells (G), granzyme B^+^ CD8^+^ T cells (H), and OVA-specific CD8^+^ T cells (I) from B16-OVA tumors in mice supplemented with Efs or Efs*-sagA* harvested five days after the start of anti–PD-L1 treatment by flow cytometry. Data are pooled from two independent experiments of 7-10 mice per group per experiment; each symbol represents one mouse. Data represent means ± s.e.m. and analyzed by Mann-Whitney U test. \**P* < 0.05, \*\**P* < 0.01, \*\*\**P* < 0.001, \*\*\*\**P* < 0.0001, ns = not significant.

Given the activity of specific enterococci during anti–PD-L1 treatment of a subcutaneous melanoma model, we subsequently interrogated whether SagA-expressing bacteria would also improve the efficacy of other checkpoint antibodies against different cancer cell types. Subcutaneous tumors were established with MCA205 fibrosarcoma or MC38 colorectal carcinoma cells in antibiotics-pretreated animals colonized by enterococci and then treated with anti–PD-1 or anti–CTLA-4, respectively. In both cases, we also observed a significant decrease in tumor growth when animals were cotreated with SagA-expressing enterococci and checkpoint inhibitor (**Fig. 3, B and C**). Given the broad efficacy of these enterococci with different targeted therapies, we then asked whether this effect was mediated by an adaptive immune response. Using our antibiotic-pretreated model, we subcutaneously implanted B16-OVA tumor cells into animals colonized with either parental *E. faecalis* or *E. faecalis–sagA* and then treated with anti–PD-L1. Tumors were harvested the day following the third antibody treatment (five days after the start of treatment), and tumor-infiltrating lymphocytes were quantified by flow cytometric analysis (**Fig. S12**). Animals colonized with *E. faecalis–sagA* showed an overall increase in the absolute amount of intratumoral CD45^+^ leukocytes as well as CD3^+^ lymphocytes (**Fig. 3, D and E**). The composition of tumor-infiltrating CD3^+^ lymphocytes showed an increase in the proportion of CD8^+^ T cells but no change in CD4^+^FoxP3^+^ regulatory T cells (**Fig. 3, F and G**). Tumors contained higher amounts of CD8^+^ T cells that expressed granzyme B (**Fig. 3H**), a marker for activated cytotoxic T lymphocytes (*24*). Tetramer staining also revealed a significant increase in the number of OVA-specific CD8^+^ T cells (**Fig. 3I**), consistent with enhanced priming of a tumor antigen-specific immune response (*9*, *13*).

To characterize the microbial mechanism of immune activation, we first examined whether SagA contributed to bacterial dissemination, as *Enterococcus* translocation from the gut has been implicated during autoimmunity (*19*), chemotherapy treatment (*25*), and alcoholic hepatitis (*26*). CFU analysis of mesenteric lymph nodes and whole spleens of supplemented animals only showed low levels of live bacteria in proximal tissues (**Fig. S10C**). The bacterial load in mesenteric lymph nodes was independent of SagA expression, suggesting that SagA did not improve barrier transit for live bacteria in our studies. We then turned our attention to peptidoglycan remodeling by SagA and the generation of muropeptides as a potential mechanism of action (*18*). Indeed, we found that Nod2, a key pattern recognition receptor for muropeptides (*27, 28*), was required for the anti–PD-L1 antitumor activity in animals supplemented with *E. faecalis–sagA* (**Fig. 4A**). To evaluate the enzymatic activity and utility of SagA to improve checkpoint blockade, we investigated heterologous expression in probiotic bacteria. *Lactococcus lactis* (Lls) has been explored extensively as a live, oral probiotic to deliver bioactive proteins and enzymes (*29*). Therefore, we produced *L. lactis* strains that chromosomally expressed wild-type, catalytically inactive (C443A), or secretion-deficient (ΔSS) SagA. All three constructs expressed well, and the wild-type and C443A mutant SagA constructs showed higher signal in the secreted fraction as expected (**Fig. 4B**). Antibiotic-pretreated animals were orally supplemented with these strains as well as parental *L. lactis* or *E. faecium* and used to monitor B16-F10 responsiveness to anti–PD-L1. Animals supplemented with *L. lactis* expressing wild-type SagA showed similar tumor growth inhibition as *E. faecium*-treated animals (**Fig. 4C**). *L. lactis* expressing catalytically inactive SagA (C443A) were not able to recapitulate the antitumor phenotype, indicating that the enzymatic activity of SagA is required to impede tumor growth. We found that the SagA secretion-deficient strain (ΔSS) did slow tumor growth, which suggests that the low amount of SagA secreted by this strain may have been sufficient to partially inhibit tumor growth. Alternatively, active, non-secreted SagA may escape from *L. lactis* due to cell lysis in the gut. Because SagA NlpC/p60 hydrolase activity was required, we asked whether synthetic muropeptide analogs of SagA enzymatic products such as muramyl dipeptide (MDP) could also elicit an improved response to checkpoint therapy. For these experiments, the Nod2-active MDP-L,D isomer or the inactive MDP-L,L diastereomer (**Fig. 4D**) were co-administered by intraperitoneal injection with anti–PD-L1. Animals that received active MDP-L,D along with anti–PD-L1 showed a significant antitumor effect that was not observed upon co-administration of the MDP-L,L negative control, suggesting the Nod2-active muropeptide agonist was sufficient to improve checkpoint blockade (**Fig. 4E**).

**Fig. 4.**
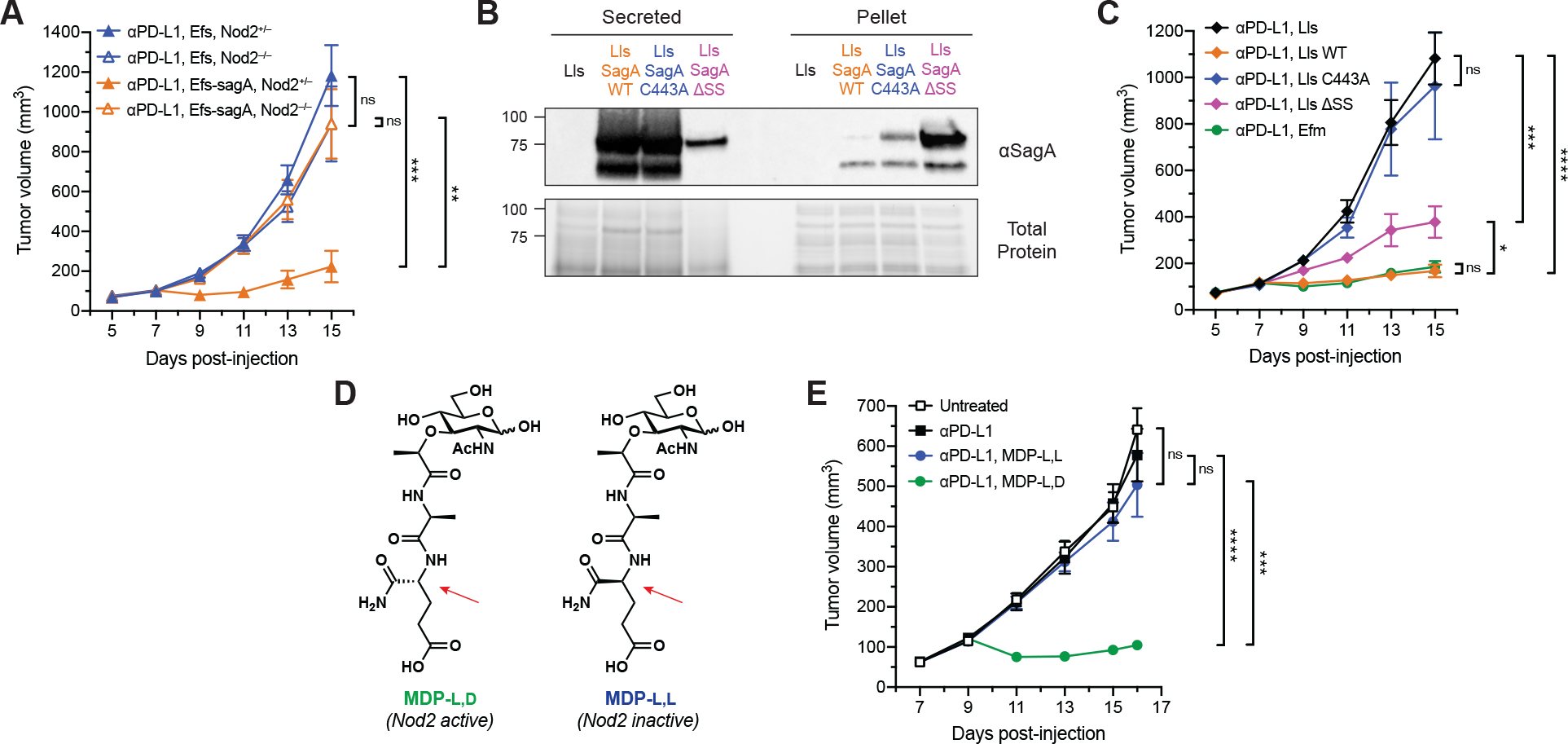
Enhancement of immunotherapy activity requires host sensing by Nod2 and peptidoglycan hydrolysis by SagA. (**A**) B16-F10 tumor growth in antibiotic-treated *Nod2^+/−^* or *Nod2^-/-^* mice that were supplemented with Efs or Efs*-sagA* and treated with anti–PD-L1 starting on day 7. *n* = 9-11 mice per group. (**B**) Western blot detection of ectopically expressed Efm Com15 SagA in secreted protein and cell pellet fractions harvested from overnight cultures of the indicated engineered *Lactococcus lactis* (Lls) strains using antiserum raised against Efm Com15 SagA. Bottom panels show total protein loading. Numbers indicate estimated molecular weight (kDa). WT = wild-type, ΔSS = signal sequence deletion. (**C**) B16-F10 tumor growth in antibiotic-treated mice that were supplemented with the indicated Lls strains and treated with anti–PD-L1 starting on day 7. *n* = 9-11 mice per group. (**D**) Chemical structures of the active (L,D) and inactive (L,L) diastereomers of muramyl dipeptide (MDP). Arrows indicate the single altered stereocenter. (**E**) B16-F10 tumor growth in antibiotic-treated, non-supplemented mice treated with anti–PD-L1 and either MDP-L,D or MDP-L,L starting on day 9. *n* = 7-8 mice per group. For (A), (C), and (E), data represent means ± s.e.m. and analyzed by mixed-effects model with Tukey’s multiple comparisons post-hoc test. *\*P* < 0.05, *\*\*P* < 0.01, \*\*\**P* < 0.001, *\*\*\*\*P* < 0.0001, ns = not significant.

Together, our data indicate that enterococci with unique NlpC/p60 peptidoglycan hydrolase activity can generate Nod2-active muropeptides and modulate the efficacy of checkpoint blockade immunotherapy *in vivo* (Fig. S13). Although bacteria enriched in responding patients do not correlate well by phylogeny (*30*), specific enterococci and other microbiota species with privileged cell wall composition and remodeling activity could provide functional indicators of therapeutic efficacy. As peptidoglycan fragments can disseminate into circulation and prime systemic immune responses (*31, 32*), NlpC/p60 hydrolases like SagA and Nod2-active muropeptides may be utilized as biomarkers to predict personalized clinical responses. Moreover, our results suggest that peptidoglycan remodeling enzymes may be utilized to reprogram probiotic bacteria and activate Nod2 during checkpoint blockade. Lastly, since MDP and Nod2 have been implicated in direct activation of macrophages for tumor clearance (*33*), epigenetic reprogramming of monocytes (*34*), generation of conventional type 1 dendritic cells (*35*), and priming of dendritic cells for cross-presentation to CD8^+^ T cells (*36*), our study highlights how microbes and small molecules that activate this pattern recognition receptor could be employed as adjuvants for cancer immunotherapy.

## Acknowledgments

We thank S. Mazel, A. Keprova, M. Jaimes, D. Tran, and the Rockefeller Flow Cytometry Resource Center for assistance with flow cytometric experiments, C. Steckler and H. Molina of the Rockefeller Proteomics Resource Center for assistance with peptidoglycan analysis, K. Eckartt for assistance in homology modeling, and M. Huse, T. Merghoub, K. Cadwell, and D. Mucida for helpful discussions.

## Author contributions

Conceptualization: M.E.G. and H.C.H.; Methodology: M.E.G., J.E., and H.C.H.; Investigation: M.E.G., J.E., and J.L.B.; Supervision: H.C.H.; Writing – original draft: M.E.G. and H.C.H.; Writing – review and editing: M.E.G., J.E., J.L.B., J.J., G.R.F., and H.C.H.; Resources – J.J. and G.R.F.

## Data and materials availability

All data are available in the main text and supplementary materials. *Enterococcus* strains used in the research are available from H.C.H. under a material transfer agreement.

## Supplementary Materials

Materials and Methods

Figures S1-S12

Tables S1-S12

References (*37–57*)

## References and Notes

1. U. Sahin, Ö. Türeci, Personalized vaccines for cancer immunotherapy. Science. 359, 1355–1360 (2018).

2. C. H. June, R. S. O’Connor, O. U. Kawalekar, S. Ghassemi, M. C. Milone, CAR T cell immunotherapy for human cancer. Science. 359, 1361–1365 (2018).

3. A. Ribas, J. D. Wolchok, Cancer immunotherapy using checkpoint blockade. Science. 359, 1350–1355 (2018).

4. E. M. Van Allen, D. Miao, B. Schilling, S. A. Shukla, C. Blank, L. Zimmer, A. Sucker, U. Hillen, M. H. G. Foppen, S. M. Goldinger, J. Utikal, J. C. Hassel, B. Weide, K. C. Kaehler, C. Loquai, P. Mohr, R. Gutzmer, R. Dummer, S. Gabriel, C. J. Wu, D. Schadendorf, L. A. Garraway, Genomic correlates of response to CTLA-4 blockade in metastatic melanoma. Science. 350, 207–211 (2015).

5. M. Łuksza, N. Riaz, V. Makarov, V. P. Balachandran, M. D. Hellmann, A. Solovyov, N. A. Rizvi, T. Merghoub, A. J. Levine, T. A. Chan, J. D. Wolchok, B. D. Greenbaum, A neoantigen fitness model predicts tumour response to checkpoint blockade immunotherapy. Nature. 551, 517–520 (2017).

6. S. Spranger, R. Bao, T. F. Gajewski, Melanoma-intrinsic β-catenin signalling prevents antitumour immunity. Nature. 523, 231–235 (2015).

7. M. Binnewies, E. W. Roberts, K. Kersten, V. Chan, D. F. Fearon, M. Merad, L. M. Coussens, D. I. Gabrilovich, S. Ostrand-Rosenberg, C. C. Hedrick, R. H. Vonderheide, M. J. Pittet, R. K. Jain, W. Zou, T. K. Howcroft, E. C. Woodhouse, R. A. Weinberg, M. F. Krummel, Understanding the tumor immune microenvironment (TIME) for effective therapy. Nat. Med. 24, 541–550 (2018).

8. M. Vétizou, J. M. Pitt, R. Daillère, P. Lepage, N. Waldschmitt, C. Flament, S. Rusakiewicz, B. Routy, M. P. Roberti, C. P. M. Duong, V. Poirier-Colame, A. Roux, S. Becharef, S. Formenti, E. Golden, S. Cording, G. Eberl, A. Schlitzer, F. Ginhoux, S. Mani, T. Yamazaki, N. Jacquelot, D. P. Enot, M. Bérard, J. Nigou, P. Opolon, A. Eggermont, P.-L. Woerther, E. Chachaty, N. Chaput, C. Robert, C. Mateus, G. Kroemer, D. Raoult, I. G. Boneca, F. Carbonnel, M. Chamaillard, L. Zitvogel, Anticancer immunotherapy by CTLA-4 blockade relies on the gut microbiota. Science. 350, 1079–1084 (2015).

9. A. Sivan, L. Corrales, N. Hubert, J. B. Williams, K. Aquino-Michaels, Z. M. Earley, F. W. Benyamin, Y. M. Lei, B. Jabri, M.-L. Alegre, E. B. Chang, T. F. Gajewski, Commensal Bifidobacterium promotes antitumor immunity and facilitates anti-PD-L1 efficacy. Science. 350, 1084–1089 (2015).

10. A. E. Frankel, L. A. Coughlin, J. Kim, T. W. Froehlich, Y. Xie, E. P. Frenkel, A. Y. Koh, Metagenomic shotgun sequencing and unbiased metabolomic profiling identify specific human gut microbiota and metabolites associated with immune checkpoint therapy efficacy in melanoma patients. Neoplasia. 19, 848–855 (2017).

11. V. Gopalakrishnan, C. N. Spencer, L. Nezi, A. Reuben, M. C. Andrews, T. V. Karpinets, P. A. Prieto, D. Vicente, K. Hoffman, S. C. Wei, A. P. Cogdill, L. Zhao, C. W. Hudgens, D. S. Hutchinson, T. Manzo, M. Petaccia de Macedo, T. Cotechini, T. Kumar, W. S. Chen, S. M. Reddy, R. Szczepaniak Sloane, J. Galloway-Pena, H. Jiang, P. L. Chen, E. J. Shpall, K. Rezvani, A. M. Alousi, R. F. Chemaly, S. Shelburne, L. M. Vence, P. C. Okhuysen, V. B. Jensen, A. G. Swennes, F. McAllister, E. Marcelo Riquelme Sanchez, Y. Zhang, E. Le Chatelier, L. Zitvogel, N. Pons, J. L. Austin-Breneman, L. E. Haydu, E. M. Burton, J. M. Gardner, E. Sirmans, J. Hu, A. J. Lazar, T. Tsujikawa, A. Diab, H. Tawbi, I. C. Glitza, W. J. Hwu, S. P. Patel, S. E. Woodman, R. N. Amaria, M. A. Davies, J. E. Gershenwald, P. Hwu, J. E. Lee, J. Zhang, L. M. Coussens, Z. A. Cooper, P. A. Futreal, C. R. Daniel, N. J. Ajami, J. F. Petrosino, M. T. Tetzlaff, P. Sharma, J. P. Allison, R. R. Jenq, J. A. Wargo, Gut microbiome modulates response to anti–PD-1 immunotherapy in melanoma patients. Science. 359, 97–103 (2018).

12. B. Routy, E. Le Chatelier, L. Derosa, C. P. M. Duong, M. T. Alou, R. Daillère, A. Fluckiger, M. Messaoudene, C. Rauber, M. P. Roberti, M. Fidelle, C. Flament, V. Poirier-Colame, P. Opolon, C. Klein, K. Iribarren, L. Mondragón, N. Jacquelot, B. Qu, G. Ferrere, C. Clémenson, L. Mezquita, J. R. Masip, C. Naltet, S. Brosseau, C. Kaderbhai, C. Richard, H. Rizvi, F. Levenez, N. Galleron, B. Quinquis, N. Pons, B. Ryffel, V. Minard-Colin, P. Gonin, J.-C. Soria, E. Deutsch, Y. Loriot, F. Ghiringhelli, G. Zalcman, F. Goldwasser, B. Escudier, M. D. Hellmann, A. Eggermont, D. Raoult, L. Albiges, G. Kroemer, L. Zitvogel, Gut microbiome influences efficacy of PD-1–based immunotherapy against epithelial tumors. Science. 359, 91–97 (2018).

13. V. Matson, J. Fessler, R. Bao, T. Chongsuwat, Y. Zha, M.-L. Alegre, J. J. Luke, T. F. Gajewski, The commensal microbiome is associated with anti–PD-1 efficacy in metastatic melanoma patients. Science. 359, 104–108 (2018).

14. F. Lebreton, A. L. Manson, J. T. Saavedra, T. J. Straub, A. M. Earl, M. S. Gilmore, Tracing the enterococci from paleozoic origins to the hospital. Cell. 169, 849–861.e13 (2017).

15. H. Hanchi, W. Mottawea, K. Sebei, R. Hammami, The genus Enterococcus: between probiotic potential and safety concerns-an update. Front. Microbiol. 9, 1791 (2018).

16. K. J. Rangan, V. A. Pedicord, Y.-C. Wang, B. Kim, Y. Lu, S. Shaham, D. Mucida, H. C. Hang, A secreted bacterial peptidoglycan hydrolase enhances tolerance to enteric pathogens. Science. 353, 1434–1437 (2016).

17. V. A. Pedicord, A. A. K. Lockhart, K. J. Rangan, J. W. Craig, J. Loschko, A. Rogoz, H. C. Hang, D. Mucida, Exploiting a host-commensal interaction to promote intestinal barrier function and enteric pathogen tolerance. Sci. Immunol. 1, eaai7732 (2016).

18. B. Kim, Y.-C. Wang, C. W. Hespen, J. Espinosa, J. Salje, K. J. Rangan, D. A. Oren, J. Y. Kang, V. A. Pedicord, H. C. Hang, Enterococcus faecium secreted antigen A generates muropeptides to enhance host immunity and limit bacterial pathogenesis. eLife. 8, e45343 (2019).

19. S. Manfredo Vieira, M. Hiltensperger, V. Kumar, D. Zegarra-Ruiz, C. Dehner, N. Khan, F. R. C. Costa, E. Tiniakou, T. Greiling, W. Ruff, A. Barbieri, C. Kriegel, S. S. Mehta, J. R. Knight, D. Jain, A. L. Goodman, M. A. Kriegel, Translocation of a gut pathobiont drives autoimmunity in mice and humans. Science. 359, 1156–1161 (2018).

20. C. K. Stein-Thoeringer, K. B. Nichols, A. Lazrak, M. D. Docampo, A. E. Slingerland, J. B. Slingerland, A. G. Clurman, G. Armijo, A. L. C. Gomes, Y. Shono, A. Staffas, M. Burgos da Silva, S. M. Devlin, K. A. Markey, D. Bajic, R. Pinedo, A. Tsakmaklis, E. R. Littmann, A. Pastore, Y. Taur, S. Monette, M. E. Arcila, A. J. Pickard, M. Maloy, R. J. Wright, L. A. Amoretti, E. Fontana, D. Pham, M. A. Jamal, D. Weber, A. D. Sung, D. Hashimoto, C. Scheid, J. B. Xavier, J. A. Messina, K. Romero, M. Lew, A. Bush, L. Bohannon, K. Hayasaka, Y. Hasegawa, M. J. G. T. Vehreschild, J. R. Cross, D. M. Ponce, M. A. Perales, S. A. Giralt, R. R. Jenq, T. Teshima, E. Holler, N. J. Chao, E. G. Pamer, J. U. Peled, M. R. M. van den Brink, Lactose drives Enterococcus expansion to promote graft-versus-host disease. Science. 366, 1143–1149 (2019).

21. A. Vermassen, S. Leroy, R. Talon, C. Provot, M. Popowska, M. Desvaux, Cell wall hydrolases in bacteria: insight on the diversity of cell wall amidases, glycosidases and peptidases toward peptidoglycan. Front. Microbiol. 10, 331 (2019).

22. J. A. Mohamed, F. Teng, S. R. Nallapareddy, B. E. Murray, Pleiotrophic effects of 2 Enterococcus faecalis sagA-like genes, salA and salB, which encode proteins that are antigenic during human infection, on biofilm formation and binding to collagen type i and fibronectin. J. Infect. Dis. 193, 231–240 (2006).

23. F. Teng, M. Kawalec, G. M. Weinstock, W. Hryniewicz, B. E. Murray, An Enterococcus faecium secreted antigen, SagA, exhibits broad-spectrum binding to extracellular matrix proteins and appears essential for E. faecium growth. Infect. Immun. 71, 5033–5041 (2003).

24. J. Lieberman, The ABCs of granule-mediated cytotoxicity: new weapons in the arsenal. Nat Rev Immunol. 3, 361–370 (2003).

25. R. Daillère, M. Vétizou, N. Waldschmitt, T. Yamazaki, C. Isnard, V. Poirier-Colame, C. P. M. Duong, C. Flament, P. Lepage, M. P. Roberti, B. Routy, N. Jacquelot, L. Apetoh, S. Becharef, S. Rusakiewicz, P. Langella, H. Sokol, G. Kroemer, D. Enot, A. Roux, A. Eggermont, E. Tartour, L. Johannes, P.-L. Woerther, E. Chachaty, J.-C. Soria, E. Golden, S. Formenti, M. Plebanski, M. Madondo, P. Rosenstiel, D. Raoult, V. Cattoir, I. G. Boneca, M. Chamaillard, L. Zitvogel, Enterococcus hirae and Barnesiella intestinihominis facilitate cyclophosphamide-induced therapeutic immunomodulatory effects. Immunity. 45, 931–943 (2016).

26. Y. Duan, C. Llorente, S. Lang, K. Brandl, H. Chu, L. Jiang, R. C. White, T. H. Clarke, K. Nguyen, M. Torralba, Y. Shao, J. Liu, A. Hernandez-Morales, L. Lessor, I. R. Rahman, Y. Miyamoto, M. Ly, B. Gao, W. Sun, R. Kiesel, F. Hutmacher, S. Lee, M. Ventura-Cots, F. Bosques-Padilla, E. C. Verna, J. G. Abraldes, R. S. Brown, V. Vargas, J. Altamirano, J. Caballería, D. L. Shawcross, S. B. Ho, A. Louvet, M. R. Lucey, P. Mathurin, G. Garcia-Tsao, R. Bataller, X. M. Tu, L. Eckmann, W. A. van der Donk, R. Young, T. D. Lawley, P. Stärkel, D. Pride, D. E. Fouts, B. Schnabl, Bacteriophage targeting of gut bacterium attenuates alcoholic liver disease. Nature. 575, 505–511 (2019).

27. D. J. Philpott, M. T. Sorbara, S. J. Robertson, K. Croitoru, S. E. Girardin, NOD proteins: regulators of inflammation in health and disease. Nat Rev Immunol. 14, 9–23 (2014).

28. R. Caruso, N. Warner, N. Inohara, G. Núñez, NOD1 and NOD2: signaling, host defense, and inflammatory disease. Immunity. 41, 898–908 (2014).

29. P. A. Bron, M. Kleerebezem, Lactic acid bacteria for delivery of endogenous or engineered therapeutic molecules. Front. Microbiol. 9, 1821 (2018).

30. V. Gopalakrishnan, B. A. Helmink, C. N. Spencer, A. Reuben, J. A. Wargo, The influence of the gut microbiome on cancer, immunity, and cancer immunotherapy. Cancer Cell. 33, 570–580 (2018).

31. T. B. Clarke, K. M. Davis, E. S. Lysenko, A. Y. Zhou, Y. Yu, J. N. Weiser, Recognition of peptidoglycan from the microbiota by Nod1 enhances systemic innate immunity. Nat. Med. 16, 228–231 (2010).

32. Z. Huang, J. Wang, X. Xu, H. Wang, Y. Qiao, W. C. Chu, S. Xu, L. Chai, F. Cottier, N. Pavelka, M. Oosting, L. A. B. Joosten, M. Netea, C. Y. L. Ng, K. P. Leong, P. Kundu, K.-P. Lam, S. Pettersson, Y. Wang, Antibody neutralization of microbiota-derived circulating peptidoglycan dampens inflammation and ameliorates autoimmunity. Nat Microbiol. 4, 766–773 (2019).

33. I. J. Fidler, S. Sone, W. E. Fogler, Z. L. Barnes, Eradication of spontaneous metastases and activation of alveolar macrophages by intravenous injection of liposomes containing muramyl dipeptide. Proc. Natl. Acad. Sci. U.S.A. 78, 1680–1684 (1981).

34. J. Kleinnijenhuis, J. Quintin, F. Preijers, L. A. B. Joosten, D. C. Ifrim, S. Saeed, C. Jacobs, J. van Loenhout, D. de Jong, H. G. Stunnenberg, R. J. Xavier, J. W. M. van der Meer, R. van Crevel, M. G. Netea, Bacille Calmette-Guerin induces NOD2-dependent nonspecific protection from reinfection via epigenetic reprogramming of monocytes. Proc. Natl. Acad. Sci. U.S.A. 109, 17537–17542 (2012).

35. D. Prescott, C. Maisonneuve, J. Yadav, S. J. Rubino, S. E. Girardin, D. J. Philpott, NOD2 modulates immune tolerance via the GM-CSF–dependent generation of CD103 ^+^ dendritic cells. Proc. Natl. Acad. Sci. U.S.A., 201912866 (2020).

36. J. Asano, H. Tada, N. Onai, T. Sato, Y. Horie, Y. Fujimoto, K. Fukase, A. Suzuki, T. W. Mak, T. Ohteki, Nucleotide oligomerization binding domain-like receptor signaling enhances dendritic cell-mediated cross-priming in vivo. J. Immunol. 184, 736–745 (2010).

37. K. S. Kobayashi, M. Chamaillard, Y. Ogura, O. Henegariu, N. Inohara, G. Nuñez, R. A. Flavell, Nod2-dependent regulation of innate and adaptive immunity in the intestinal tract. Science. 307, 731–734 (2005).

38. S. Kommineni, D. J. Bretl, V. Lam, R. Chakraborty, M. Hayward, P. Simpson, Y. Cao, P. Bousounis, C. J. Kristich, N. H. Salzman, Bacteriocin production augments niche competition by enterococci in the mammalian gastrointestinal tract. Nature. 526, 719–722 (2015).

39. O. De Henau, M. Rausch, D. Winkler, L. F. Campesato, C. Liu, D. H. Cymerman, S. Budhu, A. Ghosh, M. Pink, J. Tchaicha, M. Douglas, T. Tibbitts, S. Sharma, J. Proctor, N. Kosmider, K. White, H. Stern, J. Soglia, J. Adams, V. J. Palombella, K. McGovern, J. L. Kutok, J. D. Wolchok, T. Merghoub, Overcoming resistance to checkpoint blockade therapy by targeting PI3Kγ in myeloid cells. Nature. 539, 443–447 (2016).

40. D. Arndt, J. R. Grant, A. Marcu, T. Sajed, A. Pon, Y. Liang, D. S. Wishart, PHASTER: a better, faster version of the PHAST phage search tool. Nucleic Acids Res. 44, W16–21 (2016).

41. L.-T. Nguyen, H. A. Schmidt, A. von Haeseler, B. Q. Minh, IQ-TREE: A Fast and Effective Stochastic Algorithm for Estimating Maximum-Likelihood Phylogenies. Mol. Biol. Evol. 32, 268–274 (2015).

42. J. Trifinopoulos, L.-T. Nguyen, A. von Haeseler, B. Q. Minh, W-IQ-TREE: a fast online phylogenetic tool for maximum likelihood analysis. Nucleic Acids Res. 44, W232–W235 (2016).

43. D. T. Hoang, O. Chernomor, A. von Haeseler, B. Q. Minh, L. S. Vinh, UFBoot2: Improving the Ultrafast Bootstrap Approximation. Mol. Biol. Evol. 35, 518–522 (2018).

44. I. Letunic, P. Bork, Interactive Tree Of Life (iTOL) v4: recent updates and new developments. Nucleic Acids Res. 47, W256–W259 (2019).

45. X. Robert, P. Gouet, Deciphering key features in protein structures with the new ENDscript server. Nucleic Acids Res. 42, W320–324 (2014).

46. L. A. Kelley, S. Mezulis, C. M. Yates, M. N. Wass, M. J. E. Sternberg, The Phyre2 web portal for protein modeling, prediction and analysis. Nat. Protoc. 10, 845–858 (2015).

47. B. Kim, J. Espinosa, H. C. Hang, Biochemical analysis of NlpC/p60 peptidoglycan hydrolase activity. Methods Enzymol. 638, 109–127 (2020).

48. J. G. Caporaso, C. L. Lauber, W. A. Walters, D. Berg-Lyons, J. Huntley, N. Fierer, S. M. Owens, J. Betley, L. Fraser, M. Bauer, N. Gormley, J. A. Gilbert, G. Smith, R. Knight, Ultra-high-throughput microbial community analysis on the Illumina HiSeq and MiSeq platforms. ISME J. 6, 1621–1624 (2012).

49. R. C. Edgar, Search and clustering orders of magnitude faster than BLAST. Bioinformatics. 26, 2460–2461 (2010).

50. R. C. Edgar, SINTAX: a simple non-Bayesian taxonomy classifier for 16S and ITS sequences. bioRxiv (2016),, doi:10.1101/074161.

51. J. R. Cole, Q. Wang, J. A. Fish, B. Chai, D. M. McGarrell, Y. Sun, C. T. Brown, A. Porras-Alfaro, C. R. Kuske, J. M. Tiedje, Ribosomal Database Project: data and tools for high throughput rRNA analysis. Nucleic Acids Res. 42, D633–642 (2014).

52. P. J. McMurdie, S. Holmes, phyloseq: an R package for reproducible interactive analysis and graphics of microbiome census data. PLoS One. 8, e61217 (2013).

53. K. J. Leenhouts, J. Kok, G. Venema, Replacement recombination in Lactococcus lactis. J. Bacteriol. 173, 4794–4798 (1991).

54. G. Simons, M. Nijhuis, W. M. de Vos, Integration and gene replacement in the Lactococcus lactis lac operon: induction of a cryptic phospho-beta-glucosidase in LacG-deficient strains. J. Bacteriol. 175, 5168–5175 (1993).

55. Y. Sasaki, Y. Ito, T. Sasaki, ThyA as a selection marker in construction of food-grade host-vector and integration systems for Streptococcus thermophilus. Appl. Environ. Microbiol. 70, 1858–1864 (2004).

56. Q. N. Y. Wong, V. C. W. Ng, M. C. M. Lin, H.-F. Kung, D. Chan, J.-D. Huang, Efficient and seamless DNA recombineering using a thymidylate synthase A selection system in Escherichia coli. Nucleic Acids Res. 33, e59 (2005).

57. P. R. Jensen, K. Hammer, Minimal requirements for exponential growth of Lactococcus lactis. Appl. Environ. Microbiol. 59, 4363–4366 (1993).

