## Supplementary Materials for "*Enterococcus* peptidoglycan remodeling promotes immune checkpoint inhibitor therapy"

Supplementary Materials for
***Enterococcus* peptidoglycan remodeling promotes immune checkpoint
inhibitor therapy**

Matthew E. Griffin, Juliel Espinosa, Jessica L. Becker, Jyoti K. Jha, Gary R. Fanger, Howard C.
Hang

**This PDF file includes:**

Materials and Methods
Figs. S1 to S12
Captions for Tables S1 to S12
References (37-57)

**Other Supplementary Material for this manuscript includes the following:**

Tables S1 to S12 as a separate Excel file

### Materials and Methods

**Bacterial species.** The following commercial bacteria species and strains were sourced as follows: *Enterococcus faecium* NCTC 7171 (ATCC 19434), *E. faecium* DO/TX0016 (ATCC BAA-472), *E. faecalis* OG1RF (ATCC 47077), *E. faecalis* NCTC 775 (ATCC 19433), *E. faecalis* (ATCC 700802), *E. durans* 23C2 (ATCC 6056), *E. hirae* R (ATCC 8043), *E. mundtii* NCDO 2375 (ATCC 43186), and *E. gallinarum* NCDO 2313 (ATCC 51559). *E. faecium* Com15 was provided as a gift by Michael Gilmore (Harvard Medical School). All *L. lactis* strains were derived as described below. All *Enterococcus* strains and species were grown at 37 °C under ambient atmosphere in autoclaved Gibco Bacto Brain Heart Infusion medium (BHI, FisherScientific 237500) without antibiotics. For colony-forming unit (CFU) analysis, samples were plated onto BD BBL Enterococcosel agar medium (FisherScientific B12205).

**Cell culture.** The following cell lines were used and sourced as follows: B16-F10 (ATCC CRL-6475), MCA205 (Millipore Sigma SCC173), and MC-38 (Kerafast ENH204-FP). The B16-OVA cell line was provided as a gift by Taha Merghoub and Jedd Wolchok (Memorial Sloan Kettering Cancer Center). B16-F10 and MCA205 cells were cultured at 37 °C and 5% CO<sub>2</sub> in complete DMEM (ThermoFisher, 11995065) supplemented with 10% fetal bovine serum, 100 U mL<sup>-1</sup> penicillin, and 100 µg mL<sup>-1</sup> streptomycin. MC-38 cells were cultured at 37 °C and 5% CO<sub>2</sub> in the medium described above supplemented with 0.1 mM non-essential amino acids. Cells were maintained at no greater than 70% confluency and subcultured accordingly using TrypLE Express. Cells were used prior to 5-10 passages in vitro. Cells were also routinely cultured without antibiotics to ensure no bacterial infection and tested for mycoplasma using a Universal Mycoplasma Detection Kit (ATCC 30-1012K).

**Animals.** Specific pathogen-free, six- to eight-week old, male and female C57BL/6J (B6, 000664), B6(Cg)-*Tyr<sup>c-2J</sup>*/J (B6-albino, 000058), and B6.129S1-*Nod2<sup>tm1Flv</sup>*/J (*Nod2*<sup>-/-</sup>, 005763 from Flavell Lab, Yale School of Medicine) (37) mice were obtained from The Jackson Laboratory. When indicated, C57BL/6 mice were also obtained from Taconic Biosciences. Animals were housed in autoclaved caging with corncob bedding and paper enrichment on a 12-hour light/dark cycle. Animals were provided gamma-irradiated chow (LabDiet 5053) and sterile drinking water ad libitum. *Nod2*<sup>+/-</sup> and *Nod2*<sup>-/-</sup> littermate cohorts were generated by in-house breeding. Genotyping was performed according to the protocols established for the respective strains by The Jackson Laboratory. Animal care and experimentation were conducted in accordance with NIH guidelines and approved by the Institutional Animal Care and Use Committee at The Rockefeller University (Protocol 18048-H).

**Antibiotic pre-treatment.** Animals were provided with sterile-filtered drinking water for two weeks prior to bacterial administration supplemented with 5 g L<sup>-1</sup> streptomycin, 1 g L<sup>-1</sup> colistin sulfate, and 1 g L<sup>-1</sup> ampicillin as previously described (12). Antibiotics were dissolved in autoclaved drinking water, sterile filtered using a 0.4-micron filter, and aliquoted into two sterile 50-mL conical tubes per cage. Tubes were fitted with autoclaved #6 sipper tubes and placed onto sterile Allentown-style wire racks. Antibiotic solutions were changed at least twice weekly, and sipper tubes were changed at least once weekly.

**Bacterial administration.** Bacteria was administered through the drinking water as previously described (38). On the day prior to administration, bacteria were inoculated into 4 mL of autoclaved growth medium and grown as overnight cultures. The following day, overnight cultures were used to inoculate 50 mL of growth medium at a dilution ratio of 1:50. Bacteria were grown to late logarithmic phase ($OD \sim 1$), centrifuged at $5,000 \times g$ for 10 min, and then resuspended in sterile-filtered drinking water. Bacteria were then diluted in two 50-mL aliquots per animal cage in sterile conical tubes to $10^8$ CFU $mL^{-1}$ as previously determined by dilution plating. Tubes were then fitted with autoclaved #6 sipper tubes and provided to the animals ad libitum. For *Enterococcus* strains and species, the bacterial solutions were replaced at least twice weekly. For *L. lactis* strains, solutions were replaced every other day. Supplemented drinking water was maintained throughout the remainder of the experiment.

**Tumor inoculation.** Cells were subcultured on the day prior to harvesting for tumor inoculation. Cells were harvested using TrypLE Express, washed with and resuspended in cold phosphate-buffered saline (PBS), and counted using a hemocytometer. Cells were then resuspended at 2x concentration for injection in PBS on ice and then diluted 1:1 with Matrigel matrix (Corning, growth factor reduced, 356231). Final concentrations for each cell type are as follows: B16/F10 – $10^6$ cells $mL^{-1}$; MC-38 – $3 \times 10^6$ cells $mL^{-1}$; MCA205 – $6 \times 10^6$ cells $mL^{-1}$. Cell suspensions were kept on ice. Animals were anesthetized using 3% isoflurane, and their right flank was shaved using hair clippers. Animals were subcutaneously injected with 100 $\mu L$ of the cell suspension on their mid-right flank. The final number of injected cells for each cell type are as follows: B16/F10 – $10^5$ cells; MC-38 – $3 \times 10^5$ cells; MCA205 – $6 \times 10^5$ cells. For Fig. 3A, the experiment was conducted using the same injection volume and cell number without Matrigel.

**Tumor measurements.** Once the tumors became established at roughly 25-100 $mm^3$ as indicated in each experiment, tumor volume was measured every other day. Digital calipers were used to measure the length and width of each tumor, and tumor volume was calculated as length x width<sup>2</sup> x 0.5, where the width was the smaller of the two measurements. Animals were humanely euthanized with CO<sub>2</sub> asphyxiation at any point in the experiment if they showed tumor ulceration or lesioning, tumors larger than 1.5 $cm^3$, difficulty to ambulate, or any other moribund behavior. Generally, for B16/F10 and MC-38 tumors, measurements began on day 5. Measurements for MCA205 tumors began on day 3.

**Antibody administration.** Two days following the first tumor measurement, antibodies were administered to the animals four times every other day by intraperitoneal injection in 200 $\mu L$ antibody buffer solution (BioXCell). The antibodies and amount per injection used are as follows: anti-PD-L1 (BioXCell, BP0101) – 20 or 100 $\mu g$; anti-PD-1 (BioXCell, BP0146) – 100 $\mu g$; anti-CTLA-4 (BioXCell, BP0131) – 100 $\mu g$.

**Colony forming unit (CFU) analysis.** Fecal samples were sterilely collected three days after the start of bacterial administration. Samples were weighed, resuspended in sterile PBS, homogenized by douncing with sterile pestles, serially diluted in sterile PBS, and then plated by drip assay onto selective BD BBL Enterococcosel agar plates (Fisher Scientific, B12205). For mesenteric lymph node and spleen analyses, whole organs were sterilely dissected from animals, homogenized by douncing with sterile pestles in sterile PBS, and completely plated onto selective agar plates.

Plates were incubated for 24-48 h at 37 °C under ambient atmosphere until colonies formed. Colonies were then manually counted.

**Cell isolation and flow cytometry.** Single cell suspensions from tumors were prepared as previously described (39). Dissected tumor samples weighed, minced with scissors, and incubated in RPMI supplemented with 1.67 U mL<sup>-1</sup> Liberase TM (Roche 5401119001) and 0.2 mg mL<sup>-1</sup> DNase I (Worthington Biochemical LS002006) and at 37 °C with gentle nutation. Samples were then homogenized by pipetting and passed through a 70-µm filter (Miltenyi Biotec 130-095-823) with a sterile pestle. Cells were pelleted at 300 x g for 5 min at 4 °C, resuspended in 5 mL of red blood cell lysis buffer (ThermoFisher 00-4333-57), and incubated for 5 min at room temperature. Cells were diluted with 15 mL of PBS and pelleted as above. Cells were washed an additional two times with PBS, diluted with 0.5 mL of PBS containing 1:10,000 Zombie NIR Live/Dead stain (BioLegend 423105), and incubated for 20 min at room temperature. Samples were protected from light from this point forward. Samples were washed with 3 x 1 mL of staining buffer (PBS, 0.2% FBS) and then incubated with 20 µL of staining buffer containing 0.5 µL of TruStain FcX anti-mouse CD16/32 blocking agent (BioLegend 101319) for 20 min on ice. Cells were directly incubated with 100 µL of staining buffer containing 1:1,000 SIINFEKL tetramer-APC conjugate (NIH Tetramer Core Facility) for 20 min at room temperature. Samples were washed with 3 x 1 mL of staining buffer and then fixed and permeabilized with the FoxP3/Transcription Factor Staining Buffer Set (ThermoFisher 00-5523-00) according to the manufacturer's protocol. Cells were washed with 3 x 1 mL of perm buffer and then incubated with 20 µL of perm buffer containing 10% goat serum for 20 min on ice. Cells were then diluted with 60 µL of perm buffer containing 1.33x antibody cocktail and incubated for 20 min on ice. Antibodies were procured and used as follows: anti-CD45 (APC-F750, 30-F11, BioLegend 103153, 1:600), anti-CD3 (BV785, 17A2, Biolegend 100232, 1:300), anti-CD4 (Alexa Fluor 647, GK1.5, BioLegend 100426, 1:1,000), anti-CD8α (PE-Cy7, 53-6.7, BioLegend 100721, 1:400), anti-NK1.1 (BV480, PK136, BD Biosciences 746265, 1:80), anti-FoxP3 (Alexa Fluor 532, FJK-16s, ThermoFisher 58-5773-80, 1:40), and anti-Granzyme B (PE-CF594, GB11, BD Biosciences 562462, 1:40). After staining, cells were washed with 3 x 1 mL of perm buffer and then analyzed using a 5-laser Cytex Aurora spectral flow cytometer using SpectroFlo software (Cytex Biosciences). Doublets and dead cells were excluded based on forward and side scatter profiles and live/dead staining. Where applicable, gating was determined using fluorescence minus one (FMO) controls.

**Cloning of peptidoglycan hydrolases.** *E. faecium* SagA WT and SagA C443A constructs in pET-21a(+) were used as previously produced (16). Genomic DNA from *E. durans* 23C2, *E. hirae* R, *E. mundtii* NCDO 2375, and *E. faecalis* OG1RF was purified from 5-mL overnight cultures using the E.Z.N.A. Bacterial DNA Kit (Omega Bio-Tek D3350-01) according to the manufacturer's protocol. The appropriate genes were amplified as secretion signal deletion mutants from each genomic DNA sample using Q5 Hot-Start High-Fidelity 2x master mix (New England Biolabs M0494S) according to the manufacturer's protocol with the following primer pairs and annealing temperatures:

EdsSagpET-F: tgtttaactttaagaaggagatatacatatgGCGGACGATTTTGATTCA
EdsSagpET-R: gatctcagtggtggtggtggtggtgctcgagCATACTTACTGCAAAATCAGGT
T<sub>a</sub> = 61 °C

EhsSagpET-F: tgtttaactttaagaaggagatatacatatgGCGGACGATTTTGATTCT

EhsSagpET-R: gatctcagtggtggtggtggtggtggtgctcgagCATACTTACTGCAAAATCAGGT

T<sub>a</sub> = 61 °C

EmiSagpET-F: tgtttaactttaagaaggagatatacatatgGCGGAAGATTTTGATTCTCAAATAC

EmiSagpET-R: gatctcagtggtggtggtggtggtggtgctcgagCATACTTACAGCAAAGTCAGGTG

T<sub>a</sub> = 63 °C

EfsSalApET-F: tgtttaactttaagaaggagatatacatatgGATGAATACGATACAAAGATTCAACAAC

EfsSalApET-R: gatctcagtggtggtggtggtggtggtgctcgagGCCTGGAAAACGCCATACAAC

T<sub>a</sub> = 63 °C

EfsSalBpET-F: actttaagaaggagatatacatatgGACAATGTTGATAAAAAAATTGAAGAAAAAAATCA

EfsSalBpET-R: gatctcagtggtggtggtggtggtggtgctcgagGGCTGAGTGTCTACGATTGT

T<sub>a</sub> = 63 °C

Capitalized nucleobases indicate complementarity with the genomic DNA sequence. Sequences were inserted into the pET-21a(+) plasmid (Millipore Sigma 69740) that was double digested with NdeI (New England Biolabs R0111S) and XhoI (New England Biolabs R0146S) using the NEBuilder HiFi DNA Assembly Cloning Kit (New England Biolabs E5520S) and transformed into NEB 5-alpha chemically competent *E. coli* according to the manufacturer's protocol. Transformants were grown overnight at 37 °C on lysogeny broth agar medium (LB, Fisher Scientific BP1425) supplemented with 100 µg mL<sup>-1</sup> ampicillin. Colonies were picked and verified by Sanger sequencing (Genewiz).

**Sequence analysis, phylogeny, and homology modeling.** When possible, sequences that were directly PCR amplified from genomic DNA were used. Annotated *Enterococcus* proteome sequences were obtained from UniProt. Proteomes were searched for sequences containing InterPro annotations for peptidoglycan hydrolase domains (21). Proteins were removed if they contained either InterPro annotations for phage proteins or if they were identified as a prophage sequence by the PHASTER tool (40). Proteins were then aligned by Clustal Omega and output in FASTA format. FASTA files were uploaded to the IQ-TREE tool using default settings with the auto substitution model, and 10,000 bootstrap alignments were calculated under the ultrafast setting (41–43). The resulting phylogenetic tree files were visualized using iTOL (44). Domain regions were predicted using InterPro analysis. Sequence identity was calculated using the SIM tool via ExPASy. Alignments were visualized using ESPript 3.0 (45). Structural models for the NlpC/p60 domains of SagA orthologs were generated by Phyre2 in intensive modeling mode based on a homology search of published protein structures including *E. faecium* SagA (46). Orthologs were superimposed to align secondary structural features. MatchAlign values were calculated and reported via PyMOL v2.3.4.

**Overexpression and purification of peptidoglycan hydrolases.** The production of recombinant peptidoglycan hydrolases was adapted from a previously described protocol (18, 47). pET-21a(+) plasmids containing the peptidoglycan hydrolase genes with a C-terminal His<sub>6</sub> tag were transformed into BL21-CodonPlus (DE3)-RIL *E. coli* (Agilent 230245) according to the manufacturer's protocol and maintained in LB broth supplemented with 100 µg mL<sup>-1</sup> ampicillin and 25 µg mL<sup>-1</sup> chloramphenicol. Prior to large scale expression, 5-mL overnight cultures were grown and used to inoculate 500-mL LB cultures at a dilution of 1:100. Cultures were grown at 37 °C to mid-logarithmic phase (OD 0.5), induced with 1 mM isopropyl-D-thiogalactopyranoside, and grown for an additional 2 h at 37 °C. Cells were pelleted by centrifugation at 5,250 x g for 30 min at 4 °C and resuspended in 10 mL of lysis buffer (50 mM Bis-Tris pH 7.5, 150 mM NaCl, 0.1% SDS, 1x cOmplete EDTA-free protease inhibitor cocktail (Roche 11697498001), 2.5 U mL<sup>-1</sup> Benzonase nuclease (Millipore Sigma E1014)) at room temperature. Suspensions were transferred to 50-mL conical tubes and then sonicated 5 x 3 min in a water bath sonicator with a 1-min rest on ice between each sonication. Lysates were then transferred to ultracentrifuge tubes and separated by centrifugation at 30,000 x g for 30 min at 4 °C. After pre-rinsing 0.5 mL of His60 Ni Superflow Resin (Takara, 635662) with 5 mL of PBS, clarified lysate was added to the resin and nutated for 1 h at 4 °C. The resin bed was then washed sequentially with 2 x 5 mL of 50 mM NaH<sub>2</sub>PO<sub>4</sub> pH 8.0, 300 mM NaCl containing 20 mM and then 40 mM imidazole. Proteins were eluted from the resin with 5 mL of 50 mM NaH<sub>2</sub>PO<sub>4</sub> pH 8.0, 300 mM NaCl, 300 mM imidazole and concentrated to ≤ 0.5 mL by centrifugation at 4,000 x g at 4 °C using 10 kDa MWCO Amicon Ultra-4 centrifugal filters (Millipore Sigma UFC8010). Concentrates were next transferred to 0.5-mL 10 kDa Slide-A-Lyzer MINI dialysis devices (ThermoFisher 88401) and immersed in PBS overnight at 4 °C with gentle stirring. Proteins were further purified by fast protein liquid chromatography using a Superdex 200 Increase 10/300 GL column (GE Healthcare 28990944) pre-equilibrated with PBS. Fractions containing the target protein were combined and concentrated to > 1 mg mL<sup>-1</sup> by centrifugation at 4,000 x g at 4 °C using 10 kDa MWCO Amicon Ultra-4 centrifugal filters. Protein concentration was quantified by Pierce BCA Protein Assay Kit (ThermoFisher 23225). Proteins were aliquoted, flash frozen in liquid nitrogen, and stored at -80 °C for up to one year.

**Peptidoglycan preparation.** Isolation of peptidoglycan from Gram positive bacteria was modified from a previously described protocol (18, 47). Each *Enterococcus* species was grown as a 250-mL culture in BHI to mid-logarithmic phase (OD ~0.6) and then pelleted by centrifugation at 5,250 x g for 10 min at 4 °C. Cell pellets were then resuspended in 5 mL of 100 mM Tris-HCl pH 7.5, 0.25% SDS and boiled in a water bath for 20 min. Suspensions were then washed with 6 x 10 mL of Milli-Q H<sub>2</sub>O with centrifugation to remove wash buffer. The insoluble pellets were resuspended in 5 mL of Milli-Q H<sub>2</sub>O and sonicated in a water bath sonicator for 30 min at room temperature. Suspensions were diluted with 5 mL of 100 mM Tris-HCl pH 7.5, 10 U mL<sup>-1</sup> Benzonase and incubated for 1 h at 37 °C with shaking. Next, 5 mL of 100 mM Tris-HCl pH 7.5, 50 µg mL<sup>-1</sup> trypsin was added, and the suspension was incubated for 1 h at 37 °C with shaking. Suspensions were then boiled for 15 min, pelleted by centrifugation, and washed once with Milli-Q H<sub>2</sub>O. Pellets were then resuspended in 5 mL of 1 M HCl and incubated for 4 h at 37 °C with shaking. Suspensions were then washed with 10-mL aliquots of Milli-Q H<sub>2</sub>O until the wash buffer reached pH 4-5. The resulting purified, insoluble peptidoglycan was then resuspended in 0.5 mL of Milli-Q H<sub>2</sub>O and lyophilized in pre-weighed 1.5 mL microcentrifuge tubes. Dried samples were weighed and diluted to 25 mg mL<sup>-1</sup> with Milli-Q H<sub>2</sub>O. Samples were aliquoted, flash frozen with liquid nitrogen, and stored at -20 °C. Peptidoglycan was depolymerized to soluble muropeptides as previously described (18, 47). Insoluble peptidoglycan was diluted to 4 mg mL<sup>-1</sup> in 50 mM NaH<sub>2</sub>PO<sub>4</sub> pH 4.9 and treated with 1 mg mL<sup>-1</sup> mutanolysin from *Streptomyces globisporus* (Millipore Sigma M9901) for 16 h at 37 °C with shaking. Samples were boiled for 5 min and then pelleted by centrifugation. The soluble fractions containing soluble muropeptides were removed, lyophilized, and then either used directly for enzymatic assays or processed further for HPLC-MS analysis.

**Peptidoglycan hydrolase activity assay.** *In vitro* enzymatic assays were modified from a previously described protocol (18, 47). Purified proteins were diluted to 10 μM in 50 mM Bis-Tris pH 5.5 containing 250 μg mutanolysin-treated *E. faecalis* peptidoglycan. Samples were incubated for 16 h at 37 °C with shaking. Reactions were quenched by boiling for 5 min and centrifuged at 16,000 x g for 5 min at room temperature to remove precipitated protein. Supernatants were removed, lyophilized, and processed further for LC-MS analysis.

**HPLC-MS analysis of peptidoglycan fragment composition.** Sample preparation and analysis were performed as previously described (18, 47). Lyophilized muropeptides were resuspended in 20 μL of 250 mM NaH<sub>2</sub>BO<sub>3</sub> pH 9.0. and treated with ~1 mg NaBH<sub>4</sub>. Tubes were shaken until completely dissolved and then incubated for 1 h at room temperature. Reactions were quenched by adding 10 μL of 20% H<sub>3</sub>PO<sub>4</sub> and incubating for 1 h at room temperature. Reduced samples were centrifuged at 16,000 x g for 10 min, and supernatants were transferred to LC-MS vials. Samples were analyzed using an Agilent 1200 series LC with an LTQ Orbitrap XL MS. Generally, 15 μL of each sample was injected onto an Acclaim 120 reverse-phase C<sub>18</sub> column (5 μm particle size, 2.1 x 150 mm, ThermoFisher 059144) operated at 52 °C. Samples were run at 0.2 mL min<sup>-1</sup> with the following gradient: 0-5 min, 0% B; 5-65 min, linear gradient 0-100% B; 65-80 min, 100% B (A: 100% HPLC-grade H<sub>2</sub>O, 0.1% TFA; B: 30% HPLC-grade MeOH, 70% HPLC-grade H<sub>2</sub>O, 0.1% TFA). Products were analyzed using electrospray ionization in positive mode with acquisition over the mass range of 200-2000 *m/z*. MS data was analyzed using Xcalibur 2.2 (ThermoFisher) using fragment adduct masses calculated in ChemDraw 19.0 (PerkinElmer).

**Western blotting of supernatant and cell pellet protein fractions.** For each strain or species, 1 mL of overnight culture was pelleted by centrifugation at 5,250 x g for 10 min at room temperature. Supernatants (~900 μL) were separated and precipitated by addition of 100 μL of 100% w/v trichloroacetic acid. Samples were briefly vortexed and then incubated on ice for 1 h. Samples were then pelleted by centrifugation at 21,000 x g for 5 min at 4 °C. Pellets were washed with 2 x 500 μL ice-cold acetone and allowed to air dry. Protein was then redissolved using 100 μL of 100 μM HEPES pH 7.4, 150 mM NaCl, 4% SDS, 1x Laemmli sample buffer (Bio-Rad 161-0747). Pellets were washed with 2 x 1 mL PBS to remove residual supernatant. Pellets were then resuspended in 100 μL of 100 μM HEPES pH 7.4, 150 mM NaCl, 4% SDS, 1x Laemmli sample buffer and added to screwcap 2-mL microcentrifuge tubes containing ~100 μL of 0.1 mm zirconia/silica beads (BioSpec 11079101z). Samples were homogenized for 20 s on the highest setting using a FastPrep FP120 Cell Disrupter (ThermoFisher). Supernatant and pellet samples were then boiled for 10 min. Samples (25 μL) were resolved by SDS-PAGE using 4-15% Criterion TGX Stain-Free Midi protein gels (Bio-Rad 5678084). Total protein was visualized by Stain-Free technology using a Bio-Rad ChemiDoc MP imager. Protein was then transferred onto 0.2 μm PVDF membrane using Trans-Blot Turbo transfer system (Bio-Rad 1704273). Blots were then blocked in 50 mM Tris-HCl pH 7.5, 150 mM NaCl, 0.1% Tween-20 (TBST) containing 5% non-fat milk for 1 h at room temperature with agitation. Blots were probed overnight at 4 °C with agitation using blocking buffer containing 1:50,000 dilution of Rb antiserum raised against full-length *E. faecium* SagA as generated previously (18). Membranes were washed with 3 x 5 mL TBST for 5 min at room temperature with agitation. Blots were then probed with blocking buffer containing 1:20,000 dilution of HRP-conjugated Dk anti-Rb IgG (GE Healthcare, NA934) for 1 h at room temperature with agitation. Membranes were washed again with 3 x 5 mL TBST for 5 min at room temperature with agitation. Blots were then developed using Clarity Western ECL substrate (Bio-Rad 1705060) and imaged using a Bio-Rad ChemiDoc MP imager.

**DNA extraction for 16S rRNA sequencing.** Single fecal pellets from each animal at each timepoint were deposited a PowerBead tube (0.1 mm glass beads, Qiagen 13118-50). The extraction protocol was executed using a Maxwell RSC PureFood GMO and Authentication Kit (Promega AS1600) according to the manufacturer's protocol. Fecal pellets were resuspended in 1 mL of CTAB buffer and 20 μL RNase A solution. Samples were mixed for 10 s on a Vortex-Genie 2 (Scientific Industries SI-0236) and then incubated for 5 min at 95 °C on an Eppendorf ThermoMixer F2.0 with shaking at 1500 rpm. Tubes were then clipped onto a horizontal microtube attachment (Scientific Industries SI-H524) on the Vortex-Genie 2 and vortexed for 20 min at high speed. Samples were then centrifuged using an Eppendorf 5430R centrifuge at 12,700 rpm for 10 min at 4 °C. Samples were centrifuged again at the same settings to eliminate foam, and any residual foam or particulates were removed by sterile pipetting. Tubes were then loaded onto a Promega MaxPrep Liquid Handler instrument, which transferred 300 μL of sample into a Promega Maxwell RSC 48 extraction cartridge. Upon completion, the extraction cartridge was loaded onto a Promega Maxwell RSC 48 instrument for automated DNA extraction and elution. DNA samples were eluted in 100 μL and transferred to a standard 96-well plate. DNA was quantified in CELLSTAR 96-well black microplates (Greiner 655087) using the Quant-iT dsDNA High Sensitivity Assay Kit (ThermoFisher Q33120) with a Promega GloMax plate reader.

**16S rRNA library generation, verification, quality check, pooling, and sequencing.** Library generation was conducted using the protocol from the Earth Microbiome Project (48). Resulting amplicon libraries were washed using AMPure XP magnetic beads (Beckman Coulter A63880). Library quality and size verification was performed using the DNA 1K Reagent Kit (PerkinElmer CLS760673) on a PerkinElmer LabChip GXII instrument. Library concentrations were measured as described above after DNA extraction. Library molarity was calculated based on library peak size and concentration. Libraries were then normalized to 2 nM using the Zephyr G3 NGS Workstation (PerkinElmer 133750) and pooled together using the same volume across all normalized libraries into a 1.5-mL DNA LoBind tube (Eppendorf 022431021). Pooled libraries were sequenced on an Illumina MiSeq instrument at a loading concentration of 8 pM with 10% PhiX spike-in and 250-bp, paired-end reads using the MiSeq Reagent Kit v2 (500-cycles, Illumina MS-102-2003).

**Sequencing data processing.** Demultiplexed raw reads were processed to generate an operational taxonomic unit (OTU) table using USEARCH v11.0.667 (49). Forward and reverse reads were merged using a maximum of five mismatches and a minimum sequence identity of 90% in the overlap region, a minimum overlap length of 16 bp, and a minimum merged sequence length of 300 bp. PhiX contamination was then removed, followed by quality filtering based on FASTQ quality scores with a maximum expected error of 1.0. OTU clustering was performed using usearch -cluster\_otus with default settings. Merged (pre-filter) reads were mapped to the OTU sequences to generate the OTU table. Taxonomic classification of OTU representative sequences was performed using usearch -sintax, an implementation of the SINTAX algorithm (50) using the Ribosomal Database Project Training Set v16 (51). Alpha diversity estimation and principal coordinates analysis (PCoA) were performed using the phyloseq R package (52).

***Lactococcus lactis* strain derivation.** All *Lactococcus lactis* strains were grown in BD Difco M17 broth (FisherScientific DF1856-17-4) supplemented with 20 $\mu\text{g mL}^{-1}$ thymidine without antibiotics. The SagA expression cassette was integrated using standard double homologous recombination methods adapted from previous protocols (53, 54). DNA was synthesized encoding the *E. faecium* *sagA* sequence flanked by 500 bp overhangs homologous to the upstream and downstream *thyA* target gene sequence (denoted as SagA-*thyA*). SagA-*thyA* was inserted into the multiple cloning site of the pRISE1.2 plasmid, which contains a temperature sensitive origin of replication. pRISE1.2-SagA-*thyA* was transformed into *L. lactis*, and cultures were maintained at 37 °C to initiate integration via double crossover to produce a thymidine auxotroph. A positive selection system to identify positive crossover events was adapted from previous protocols (55, 56). Transformed *L. lactis* cultures were grown overnight in minimal medium (57) containing 50 $\mu\text{g mL}^{-1}$ erythromycin to select for transformants, 1-5 $\mu\text{g mL}^{-1}$ trimethoprim to eliminate bacteria with functional *thyA*, and 20 $\text{mg mL}^{-1}$ thymidine to maintain auxotrophic growth. Cultures were diluted 1:1,000 into minimal medium with the same supplements for a second round of overnight culture. After selection, bacteria were plated on minimal medium agar plates supplemented with trimethoprim and thymidine, and individual clones were selected. Stability of the integration event was confirmed by growth the selected clones under non-selective conditions for 100 generations with 1,000-fold dilutions every 24 h for 3 days. Similar protocols were used to generate the null *thyA* auxotroph, SagA C443A mutant, and SagA $\Delta\text{SS}$ mutant.

**Statistical analyses.** Non-sequence data were analyzed with Microsoft Excel 16.35 and Prism 8.4.0 (GraphPad). Tumor growth assays were analyzed using a mixed-effects analysis on the log pre-processed tumor volumes with Tukey's multiple comparisons post hoc tests. Flow cytometric data were compared using a non-parametric Mann-Whitney U test. $P < 0.05$ was considered statistically significant for all assays, and individual $P$ values are denoted as follows: \*$P < 0.05$, \*\*$P < 0.01$, \*\*\*$P < 0.001$, \*\*\*\*$P < 0.0001$.

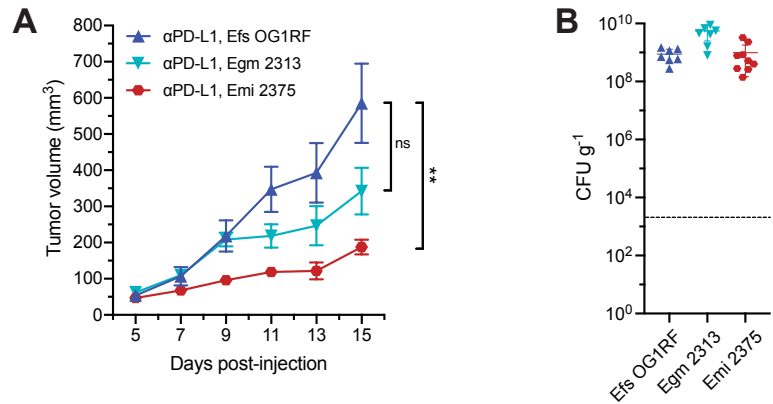

**Figure S1. Antitumor activity of *Enterococcus* species.** (A) B16-F10 tumor growth in antibiotic-treated animals that were supplemented with the indicated *Enterococcus* strains and treated with anti-PD-L1 starting on day 7 similar to Fig. 1E, $n = 7-8$ animals per group. Data points represent means ± s.e.m. and are analyzed by mixed-effects model with Tukey's multiple comparisons post hoc test. $**P < 0.01$, ns = not significant. (B) Colony forming unit (CFU) analysis of *Enterococcus* species in fecal samples harvested from supplemented animals in (A), $n = 7-8$ per group. Each symbol represents one mouse. The dotted line indicates the limit of detection (2,000 CFU g<sup>-1</sup>). Data represent means ± 95% confidence interval.

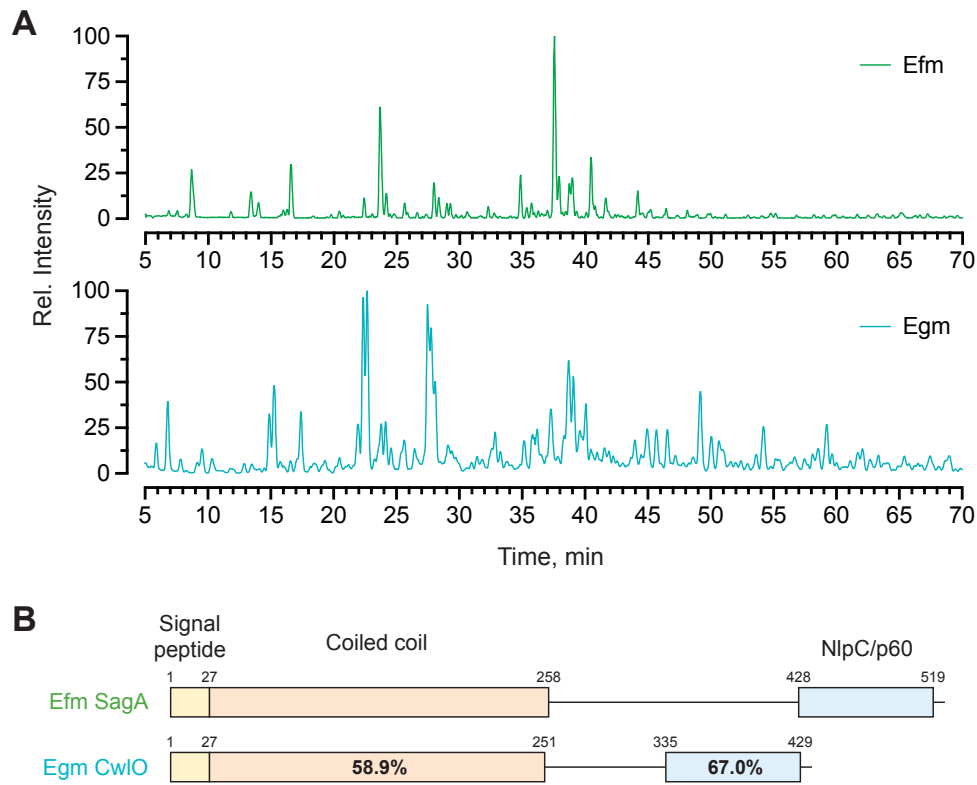

**Figure S2. Peptidoglycan profile and closest NlpC/p60 homolog are distinct in *E. gallinarum*.** (A) LC-MS ion chromatograms of mutanolysin-treated peptidoglycan harvested from *E. faecium* Com15 (*Efm*) and *E. gallinarum* 2375 (*Egm*) similar to Fig. 2A. Data are shown as relative intensity of base peak ion abundance. (B) Comparison of primary sequence homology and domain architecture of *E. faecium* Com15 SagA and the closest ortholog from *Egm* 2375 similar to Fig. 2D. Numbers above each bar denote amino acid residue boundaries of the indicated domains, and percentages represent sequence identity.

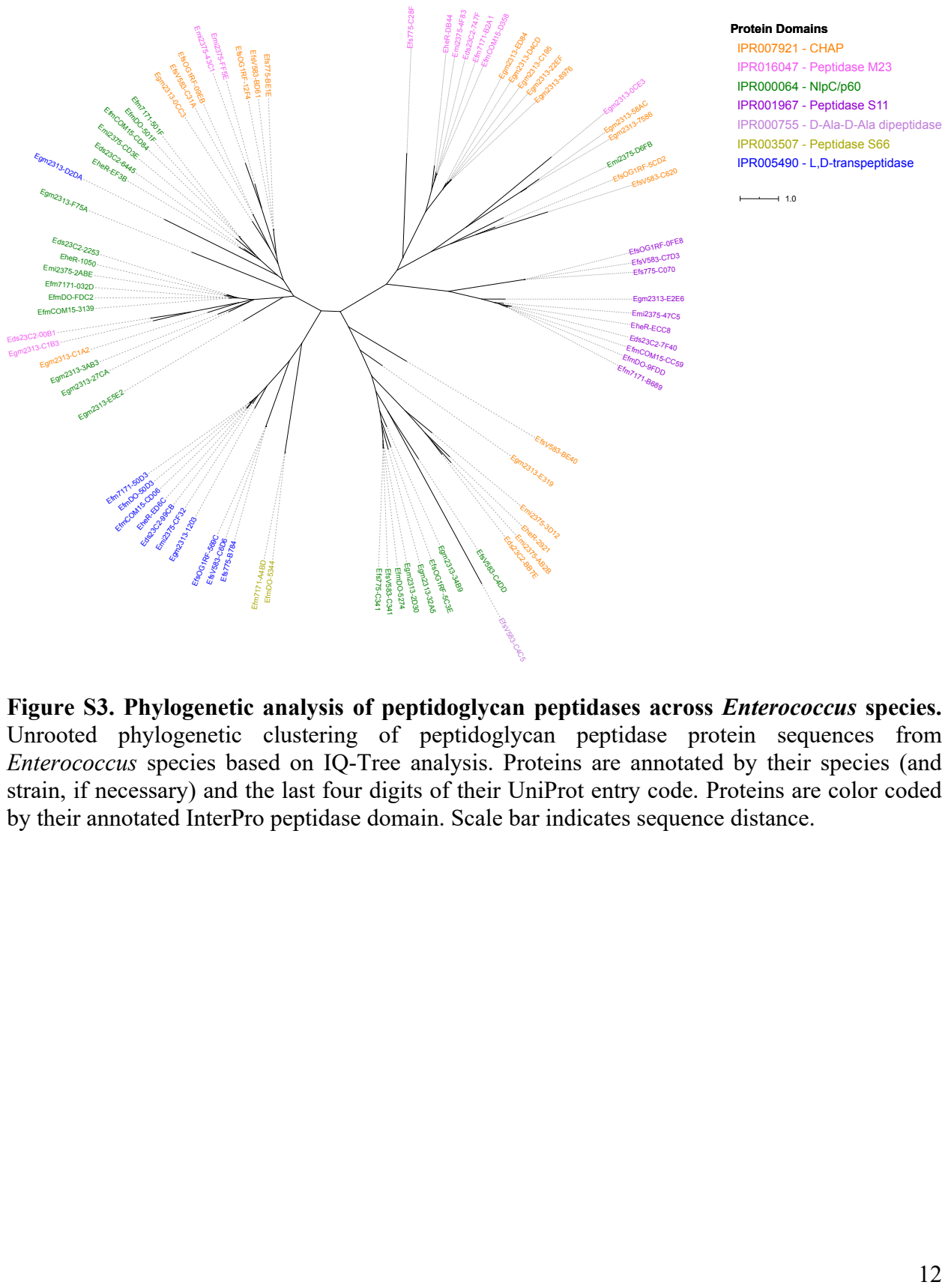

Figure S3. Phylogenetic analysis of peptidoglycan peptidases across Enterococcus species.

Unrooted phylogenetic clustering of peptidoglycan peptidase protein sequences from Enterococcus species based on IQ-Tree analysis. Proteins are annotated by their species (and strain, if necessary) and the last four digits of their UniProt entry code. Proteins are color coded by their annotated InterPro peptidase domain. Scale bar indicates sequence distance.

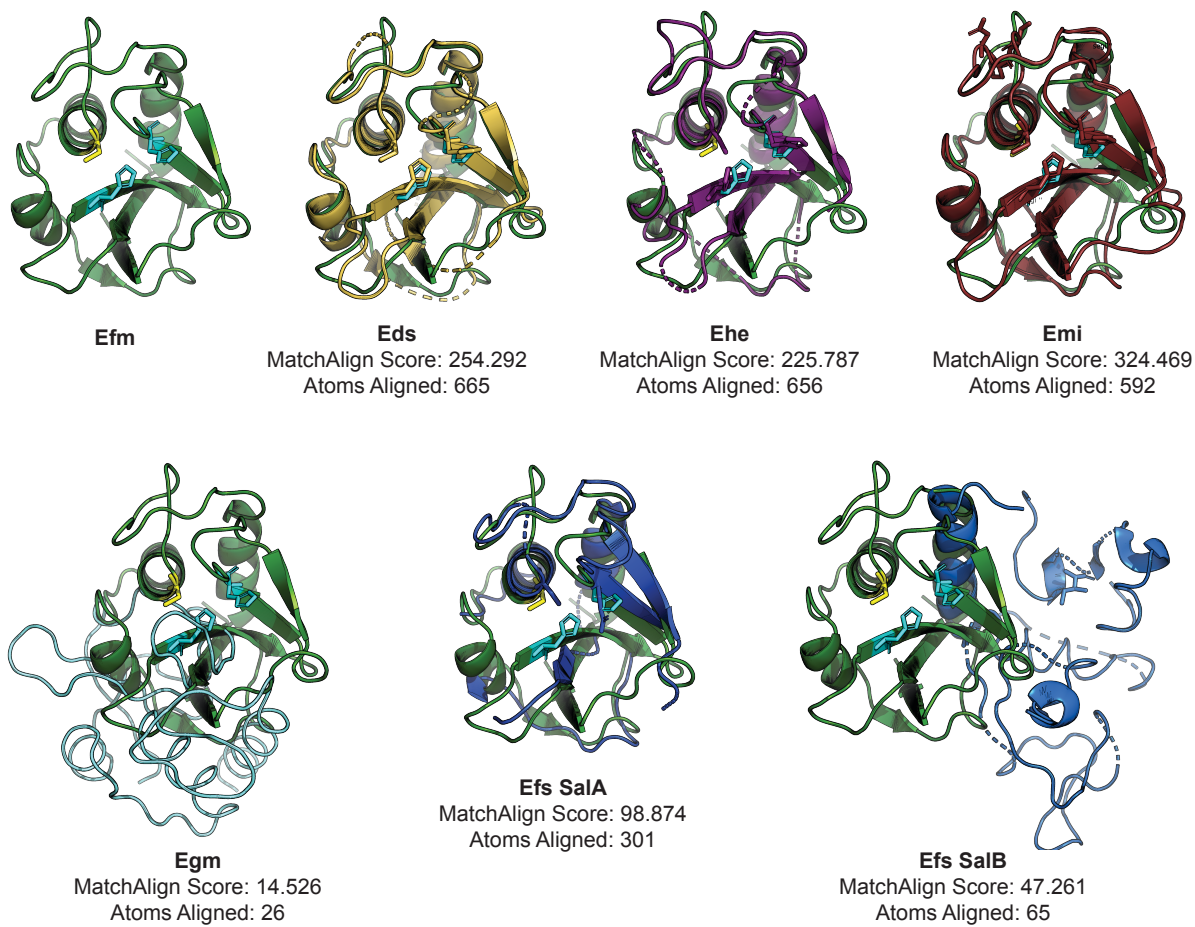

**Figure S4. Three-dimensional homology modeling of SagA orthologs.** Structural models for the catalytic domains of putative SagA orthologs from the annotated *Enterococcus* species as well as SalA and SalB from *Efs* were produced using Phyre2 using a homology search against published structures including the NlpC/p60 domain of *Efm* SagA. Structures were aligned and scored using PyMOL. The amino acid residues of the putative catalytic triad in each structure are shown using stick models.

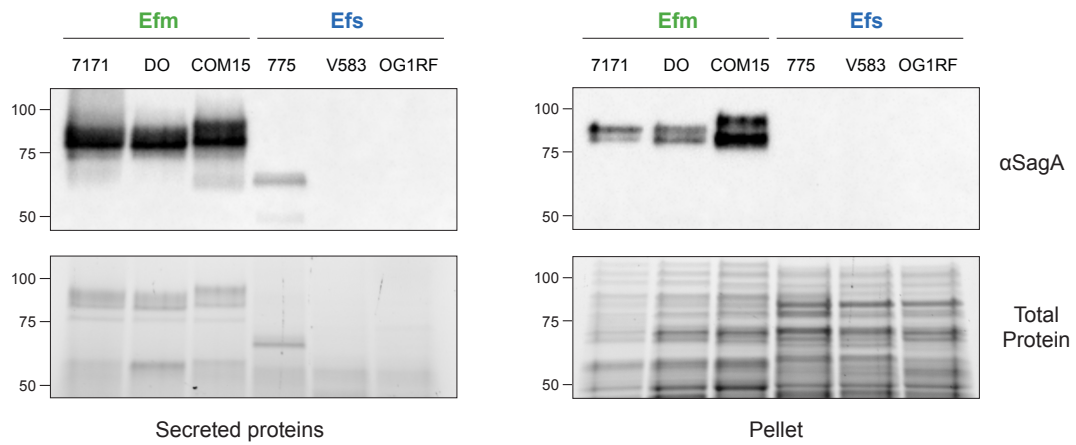

**Figure S5. Expression and secretion of SagA in *E. faecium* and *E. faecalis* strains.** Western blot detection of SagA orthologs in secreted protein and cell pellet fractions harvested from overnight cultures of the indicated *Efm* and *Efs* strains using antiserum raised against *Efm* Com15 SagA. Bottom panels show total protein loading. Numbers indicate estimated molecular weight (kDa).

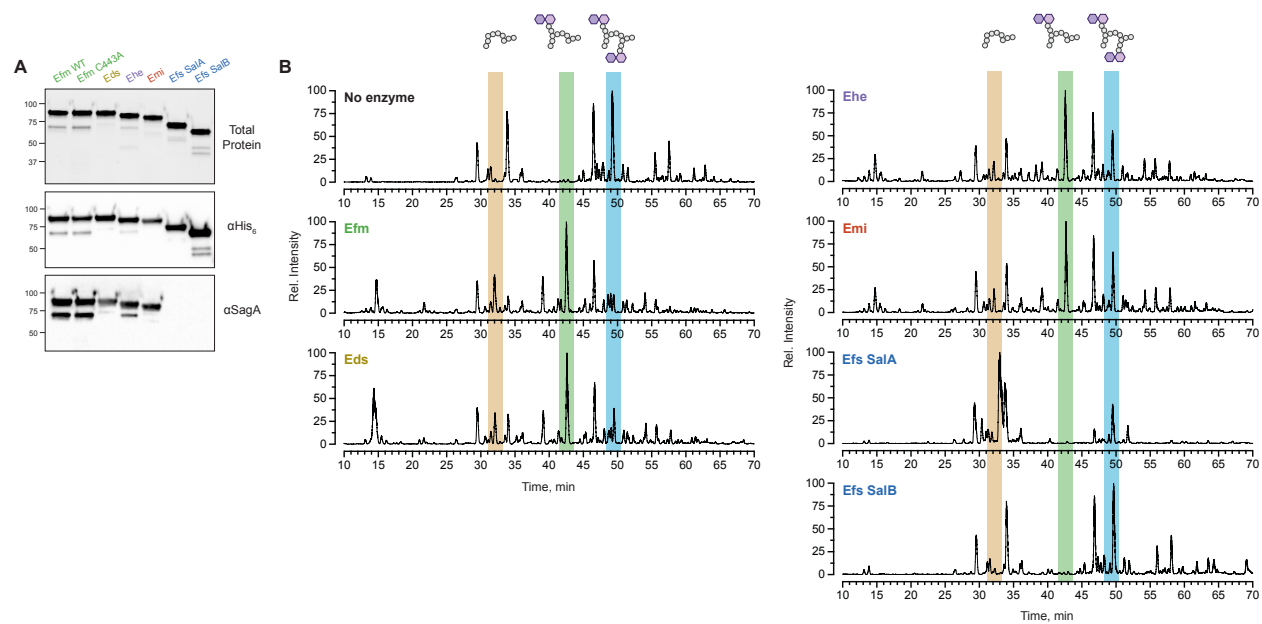

**Figure S6. Biochemical purification and characterization of SagA orthologs.** (A) Total protein and Western blotting of full-length SagA orthologs after expression and purification from *E. coli*. The two SagA-like proteins from *Efs* OG1RF, SalA and SalB, do not show reactivity using antiserum raised against *Efm* Com15 SagA. (B) In vitro activity of purified SagA orthologs corresponding to Fig. 2E. Full LC-MS ion chromatograms of a mixture of peptidoglycan fragments incubated with purified SagA orthologs from the indicated species for 16 hours at 37 °C. Data are shown as relative intensity of base peak ion abundance. The blue column indicates a crosslinked substrate for SagA that is cleaved iteratively into two products indicated by the green and orange columns, corresponding to the loss of one or two GlcNAc-muramyl dipeptide molecules, respectively. See Fig. S7 for the corresponding chemical structures.

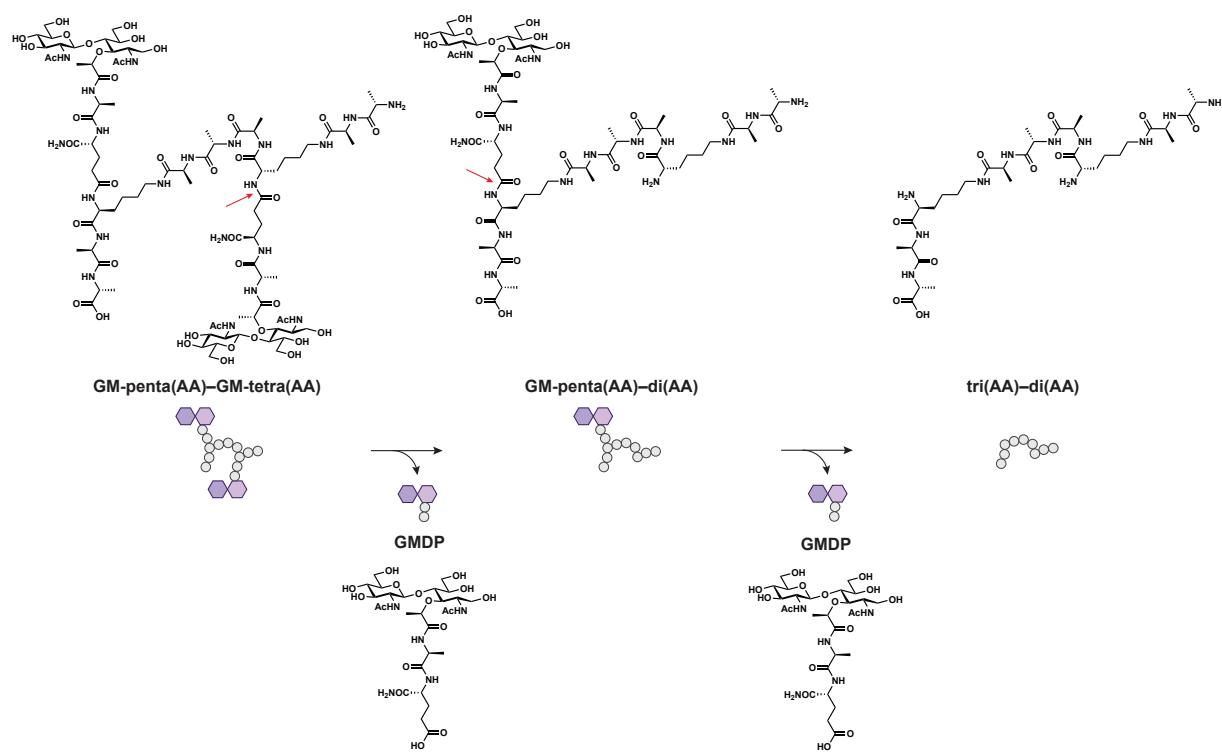

**Figure S7. Chemical structures of peptidoglycan substrate and products of SagA activity.** Chemical structures of the model crosslinked peptidoglycan substrate (GM-penta(AA)–GM-tetra(AA), blue column in LC-MS ion chromatograms) and the two products (GM-penta(AA)–di(AA), green column; tri(AA)–di(AA), orange column) produced by SagA and its orthologs through the iterative hydrolysis of D-isoglutamine–L-lysine amide bonds indicated by the red arrows. Each hydrolysis step releases one equivalent of GlcNAc-muramyl dipeptide (GMDP).

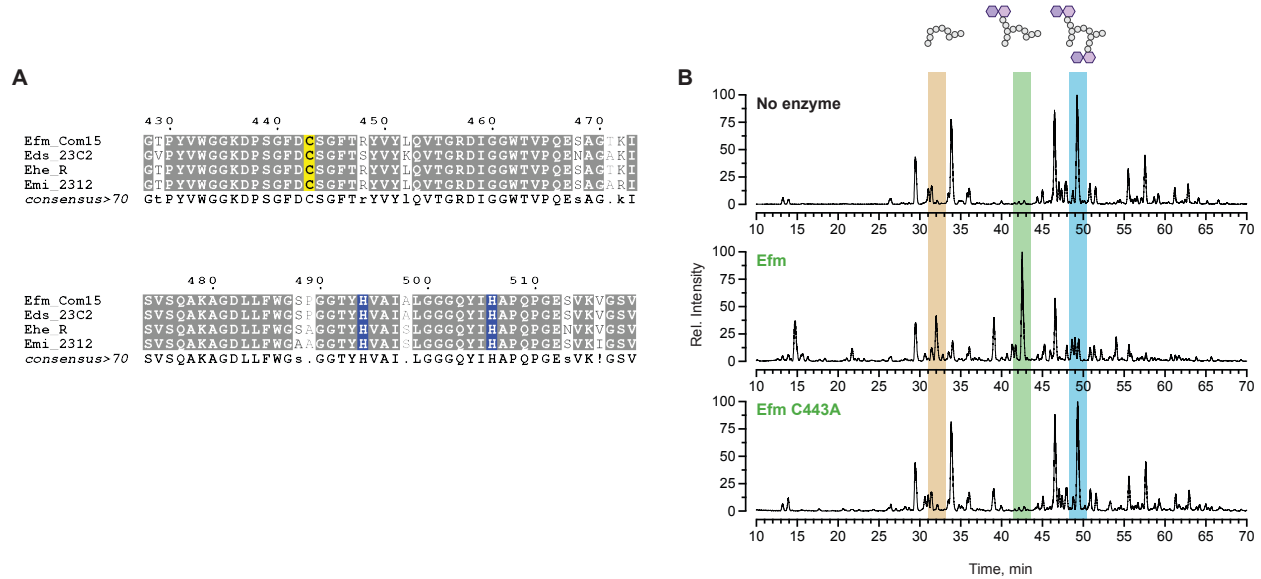

**Figure S8. SagA activity requires a conserved cysteine residue.** (A) Alignment of the NlpC/p60 domains of SagA orthologs from active *Enterococcus* species. Conserved residues are annotated in gray, and the putative catalytic triad (Cys-His-His) is highlighted in yellow and blue. (B) In vitro activity of purified SagA orthologs corresponding to Fig. 2E and Fig. S6B. Full LC-MS ion chromatograms of a mixture of peptidoglycan fragments incubated with purified *Efm* Com15 wild-type or C443 mutant SagA for 16 hours at 37 °C. Data are shown as relative intensity of base peak ion abundance. Colored columns represent peptidoglycan substrate and products as described in Fig. S6.

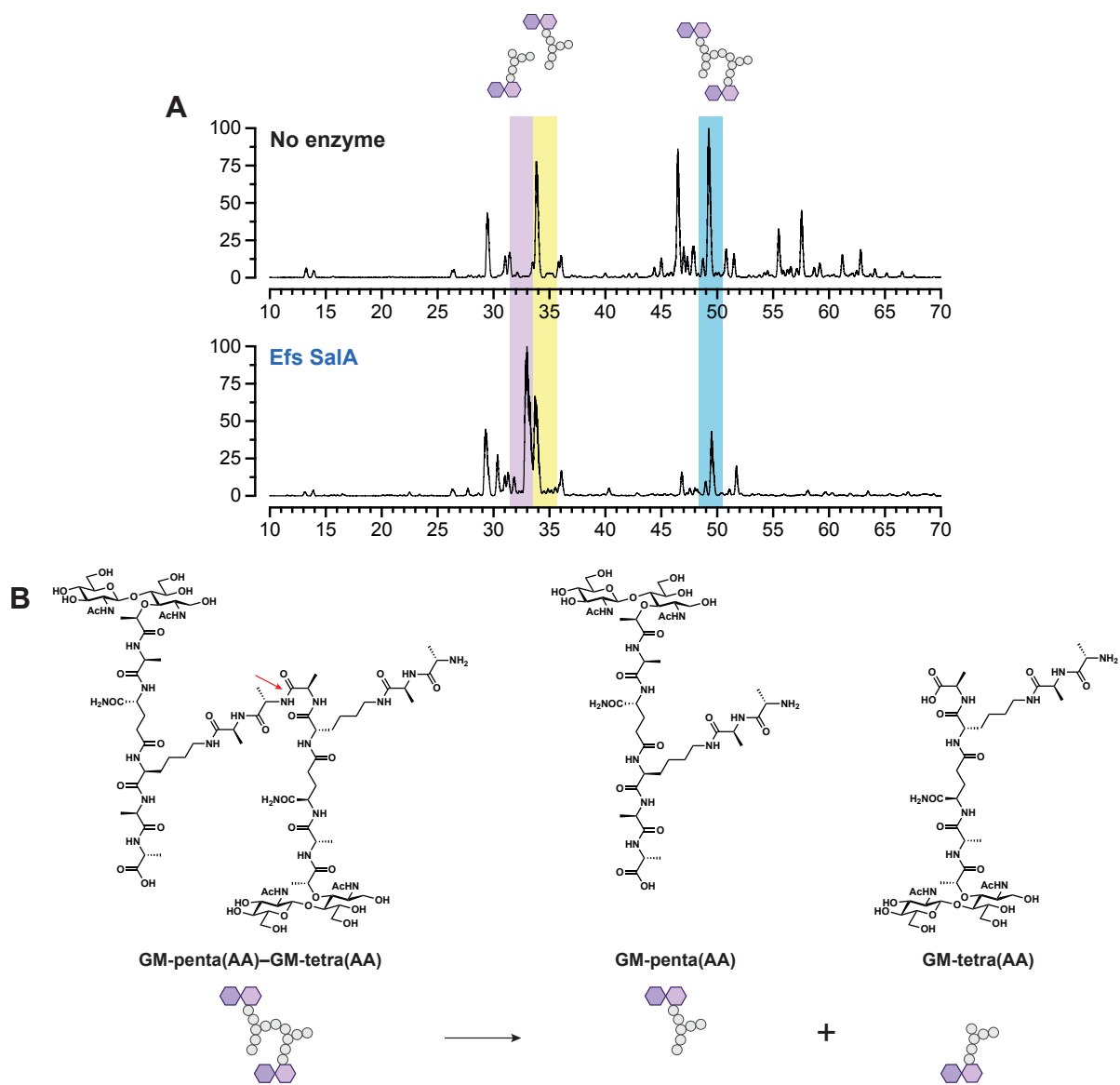

**Figure S9. *E. faecalis* SalA exhibits distinct D,L-endopeptidase activity.** (A) In vitro activity of purified *Efs* OG1RF SalA corresponding to Figs. 2E and S6B. Full LC-MS ion chromatograms of a mixture of peptidoglycan fragments incubated with purified *Efs* SalA for 16 hours at 37 °C. Data are shown as relative intensity of base peak ion abundance. The blue column indicates a SalA substrate that is cleaved at the cross-bridge into two products indicated by the yellow and purple columns. (B) Chemical structures of the model crosslinked peptidoglycan substrate (GM-penta(AA)-GM-tetra(AA), blue column in LC-MS chromatograms) and the two products (GM-penta(AA), yellow column; GM-tetra(AA), purple column) produced by *Efs* SalA.

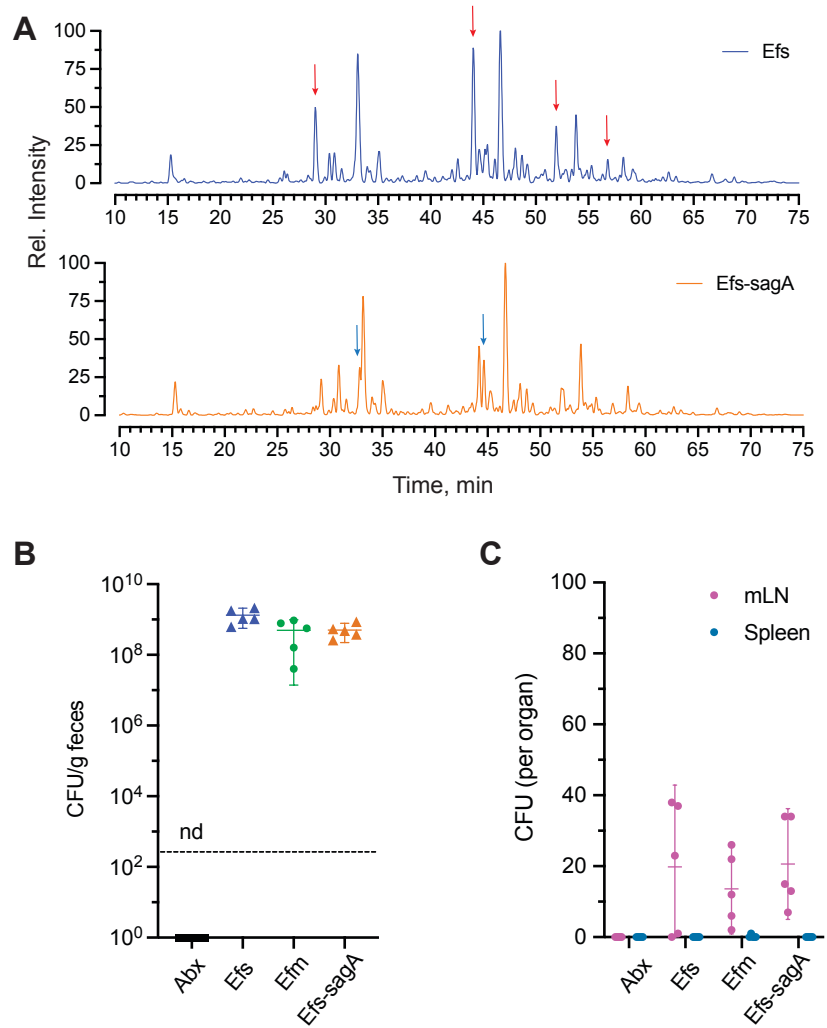

**Figure S10. In vitro and in vivo characterization of *Efs-sagA*.** (A) LC-MS ion chromatograms of mutanolysin-treated peptidoglycan harvested from parental *Efs* and *Efs-sagA* similar to Fig. 2A. Data are shown as relative intensity of base peak ion abundance. Red arrows indicate SagA substrate fragments found in the parental *Efs* profile that decrease in the *Efs-sagA* profile. Blue arrows indicate SagA product fragments that increase in the *Efs-sagA* profile. (B) Colony forming unit (CFU) analysis of *Enterococcus* species in fecal samples harvested from supplemented animals similar to Fig. 1F, $n = 5$ per group. The dotted line indicates the limit of detection (2,000 CFU g<sup>-1</sup>). nd = not detected. (C) CFU analysis of disseminated *Enterococcus* species in mesenteric lymph nodes and spleens harvested from supplemented animals in (B), $n = 5$ per group. For (B) and (C), each symbol represents one mouse. Data represent means ± 95% confidence interval.

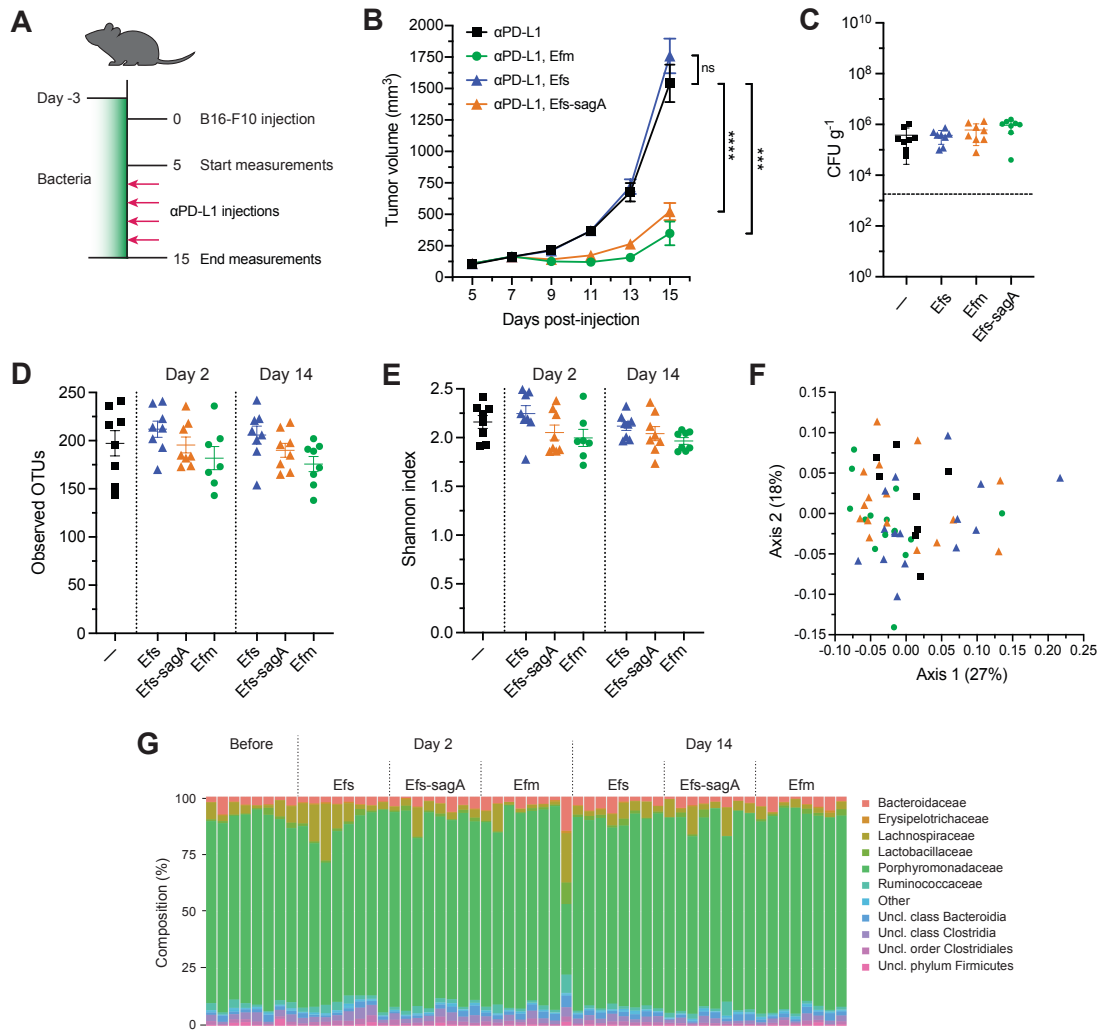

**Figure S11. *Efm* and *Efs-sagA* potentiate checkpoint blockade in an intact microbiota.** (A) Schematic of tumor growth model in Taconic mice with oral *Enterococcus* supplementation. The model was identical to that presented in Fig. 1A except that Taconic mice were used and no antibiotics were given to animals prior to supplementation. (B) B16-F10 tumor growth in Taconic animals that were supplemented with the indicated *Enterococcus* species and treated with anti-PD-L1 starting on day 7, $n = 8$ animals per group. Data represent means ± s.e.m. and were analyzed by mixed-effects model with Tukey's multiple comparisons post hoc test; $**P < 0.01$, ns = not significant. (C) Colony forming unit (CFU) analysis of *Enterococcus* species in fecal samples harvested from supplemented animals in (B), $n = 8$ per group. The dotted line indicates the limit of detection (2,000 CFU g<sup>-1</sup>). Data represent means ± 95% confidence interval. (D) Observed operational taxonomic units (OTUs) measured by 16S rRNA sequencing from fecal samples collected from animals in (B) prior to supplementation or 2 or 14 days after the start of supplementation, $n = 7-8$ animals per group. (E) Shannon diversity indices, (F) principal coordinates analysis of Bray-Curtis similarity values, and (G) family-level taxonomic compositions for samples described in (D). Uncl. = unclassified. For (C) to (G), each symbol or bar represents one animal. For (D), (E), and (F), samples with less than 10,000 reads ($n = 1$; *Efm* Day 2, #8) were excluded from analysis.

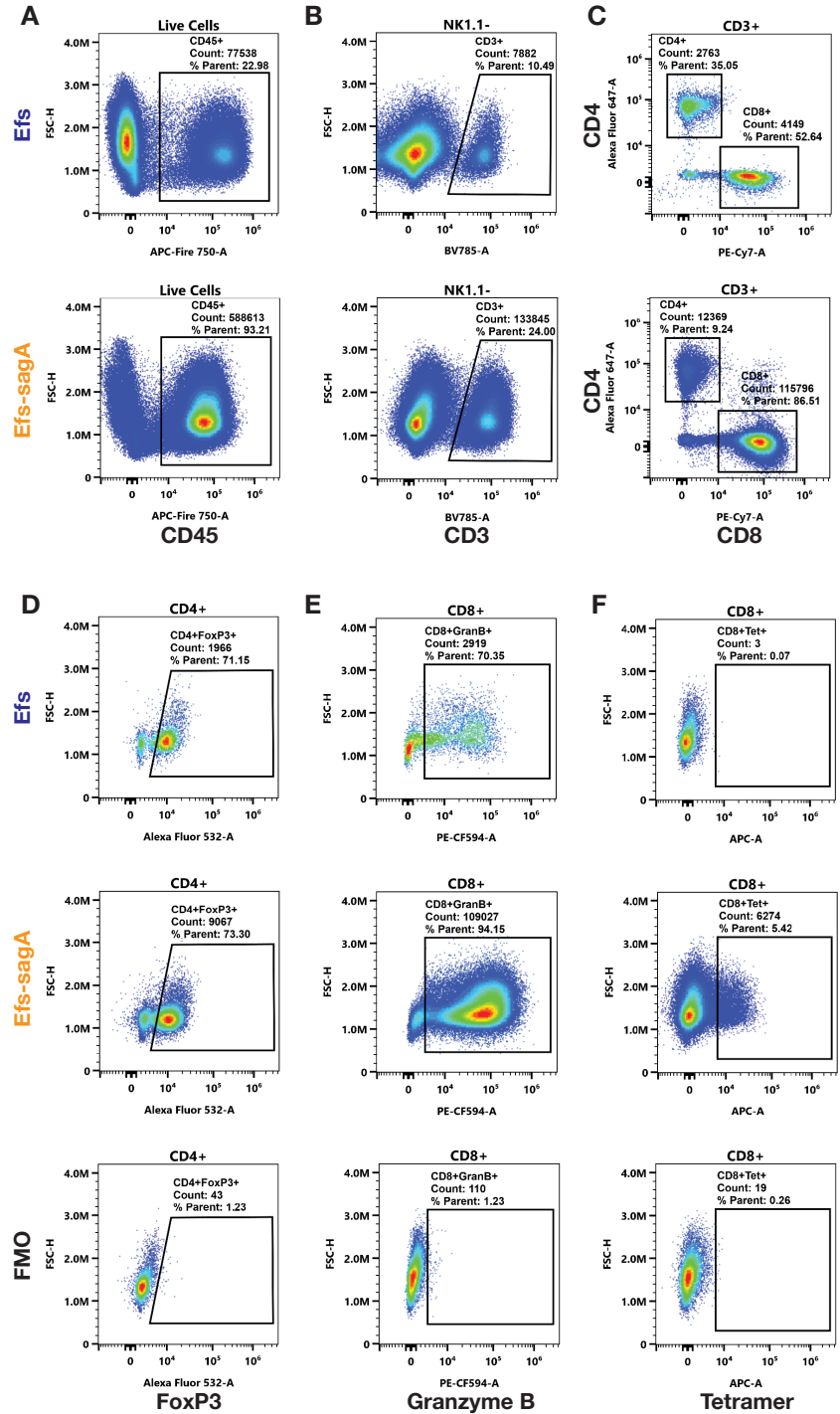

**Figure S12. Flow cytometry plots of tumor-infiltrating lymphocytes.** Representative plots of single cell suspensions harvested from B16-OVA tumors in animals colonized with *E. faecalis* or *E. faecalis-saga* corresponding to Fig. 3, D-I. Graphs depict (A) total live CD45<sup>+</sup> cells, (B) CD45<sup>+</sup>NK1.1<sup>-</sup>CD3<sup>+</sup> T lymphocytes, (C) CD4 and CD8 compartments of CD45<sup>+</sup>CD3<sup>+</sup> T lymphocytes, (D) FoxP3<sup>+</sup>CD4<sup>+</sup> regulatory T cells, (E) GranzymeB<sup>+</sup>CD8<sup>+</sup> cytotoxic T cells, and (F) tetramer<sup>+</sup>CD8<sup>+</sup> tumor OVA antigen-specific T cells. For (D)-(F), fluorescence minus one (FMO) controls used to define gates are included.

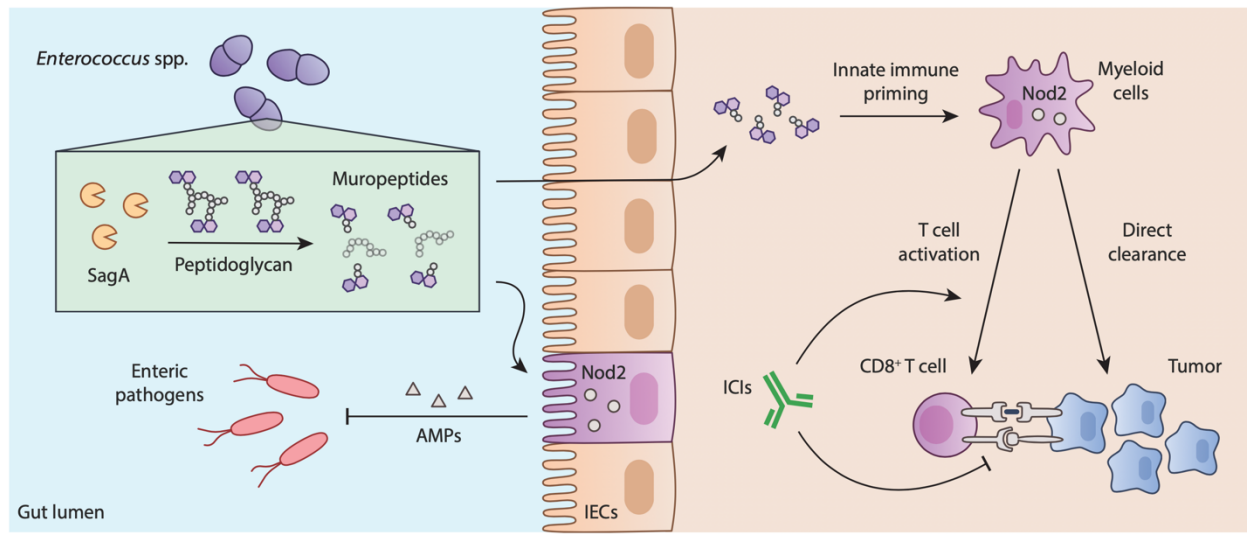

**Figure S13. Active enterococci improve immune checkpoint inhibitor efficacy.** Some *Enterococcus* species express orthologs of the NlpC/p60 peptidoglycan hydrolase SagA, which can remodel peptidoglycan to produce muropeptide fragments. The generation of these immune active metabolites can improve tolerance toward enteric pathogens or increase checkpoint blockade immunotherapy in a Nod2-dependent manner.

**Additional data tables S1 through S12 are included in a separate Excel (.xlsx) file.**

**Table S1.** Domain names and corresponding InterPro (IPR) IDs used to search the proteomes of *Enterococcus* strains and species for peptidoglycan remodeling peptidases.

**Table S2.** UniProt proteome IDs, NCBI organism IDs, and alternative designations of *Enterococcus* strains and species used herein.

**Tables S3-S12.** Putative peptidoglycan remodeling peptidases of *Efm* Com15 (S3), *Efm* 7171 (S4), *Efm* DO (S5), *Eds* 23C2 (S6), *Ehe* R (S7), *Emi* 2375 (S8), *Efs* OG1RF (S9), *Efs* 775 (S10), *Efs* V583 (S11), and *Egm* 2313 (S12). Each table contains the UniProt proteome entry ID, alternative entry ID(s), gene and protein names (if annotated), InterPro domain IDs, and primary sequence of every protein with a matching InterPro domain ID from Table S1 within the corresponding UniProt proteomes from Table S2.
